# Programmable RNA recognition using a CRISPR-associated Argonaute

**DOI:** 10.1101/208041

**Authors:** Audrone Lapinaite, Jennifer A. Doudna, Jamie H. D. Cate

**Affiliations:** Department of Molecular and Cell Biology, University of California, Berkeley, CA 94720; Molecular Biophysics and Integrated Bioimaging Division, Lawrence Berkeley National Laboratory, Berkeley, California 94720; Howard Hughes Medical Institute, University of California, Berkeley, CA 94720; Innovative Genomics Institute, University of California, Berkeley, CA 94720; and; Department of Chemistry, University of California, Berkeley, CA 94720

**Keywords:** Argonaute, CRISPR, programmable RNA recognition, small noncoding RNA, RNA editing, inosine

## Abstract

Argonaute proteins (Agos) are present in all domains of life. While the physiological function of eukaryotic Agos in regulating gene expression is well documented, the biological roles of many of their prokaryotic counterparts remain enigmatic. In some bacteria, Agos are associated with CRISPR (Clustered Regularly Interspaced Short Palindromic Repeats) loci and use non-canonical 5’-hydroxyled guide RNAs (gRNAs) for nucleic acid targeting. Here we show that using 5-bromo-2′-deoxyuridine (BrdU) as the 5’ nucleotide of gRNAs stabilizes *in vitro* reconstituted CRISPR-associated *Marinitoga piezophila* Argonaute-gRNA complexes (MpAgo RNPs) and significantly improves their specificity and affinity for RNA targets. Using reconstituted MpAgo RNPs with 5’-BrdU modified gRNAs, we mapped the seed region of the gRNA, and identified the nucleotides of the gRNA that play the most significant role in targeting specificity. We also show that these MpAgo RNPs can be programmed to distinguish between substrates that differ by a single nucleotide, using permutations at the 6^th^ and 7^th^ positions in the gRNA. Using these specificity features, we employed MpAgo RNPs to detect specific Adenosine to Inosine edited RNAs in a complex mixture. These findings broaden our mechanistic understanding of the interactions of Argonautes with guide and substrate RNAs, and demonstrate that MpAgo RNPs with 5’-BrdU modified gRNAs can be used as a highly-specific RNA-targeting platform to probe RNA biology.

**SIGNIFICANCE:** Argonaute proteins are present in bacteria, archaea and eukaryotes. They play an important role in a wide range of biological processes, from transcriptional and translational gene expression regulation to defense against viruses and silencing of mobile genetic elements. Here we present mechanistic insights into the interactions of the CRISPR-associated *Marinitoga piezophila* Argonaute (MpAgo) with its guide RNA (gRNA) and RNA substrates. By modifying the 5’-nucleotide of the gRNA, we demonstrate that MpAgo-gRNA complexes (RNPs) are easily programmable, have high affinity to fully complementary RNA substrates, and can discriminate by over 300 fold between substrates that differ by only a single nucleotide. These MpAgo RNPs should be useful for probing endogenous RNAs in living cells.

## INTRODUCTION

Argonautes are nucleic acid-guided proteins present in organisms from all three domains of life (1). In eukaryotes, Argonautes (eAgos) play central roles in RNA interference (RNAi) and micro-RNA (miRNA) pathways that coordinate a wide range of cellular processes including transcriptional and translational gene regulation (2, 3), silencing of mobile genetic elements (4, 5), and host defense (6). eAgos bind single-stranded RNAs (ssRNA), either small interfering RNAs (siRNAs), miRNAs or Piwi-interacting RNAs (piRNAs), which act as templates for the recognition of complementary ssRNA targets, leading either to Ago cleavage of the targeted RNA (7) or to the recruitment of additional components of the RNA degradation machinery (9–11).

Although prokaryotes lack RNAi pathways (12), prokaryotic Argonautes (pAgos) are thought to contribute to host defense against foreign DNA (13). Despite their structural similarity to eAgos (Fig. 1A), pAgos are more diverse in their biochemical behavior than their eukaryotic counterparts. For example, some pAgos can use either DNA or RNA guides, and can target either DNA or RNA substrates (8, 13–15). Differing from other Agos, the CRISPR locus-associated pAgo subfamily found in *Marinitoga piezophila, Thermotoga profunda* and *Marinitoga sp. 1155)* (16) uses 5’-hydroxylated (5’-OH) guide RNAs (gRNAs) rather than 5’-phosphorylated guides. Furthermore, the *Marinitoga piezophila* Ago (MpAgo) cleaves both ssDNA and ssRNA *in vitro* in a gRNA-dependent manner (16). However, the origin of endogenous MpAgo guides and the physiological function of MpAgo remain enigmatic.

**Figure 1.**
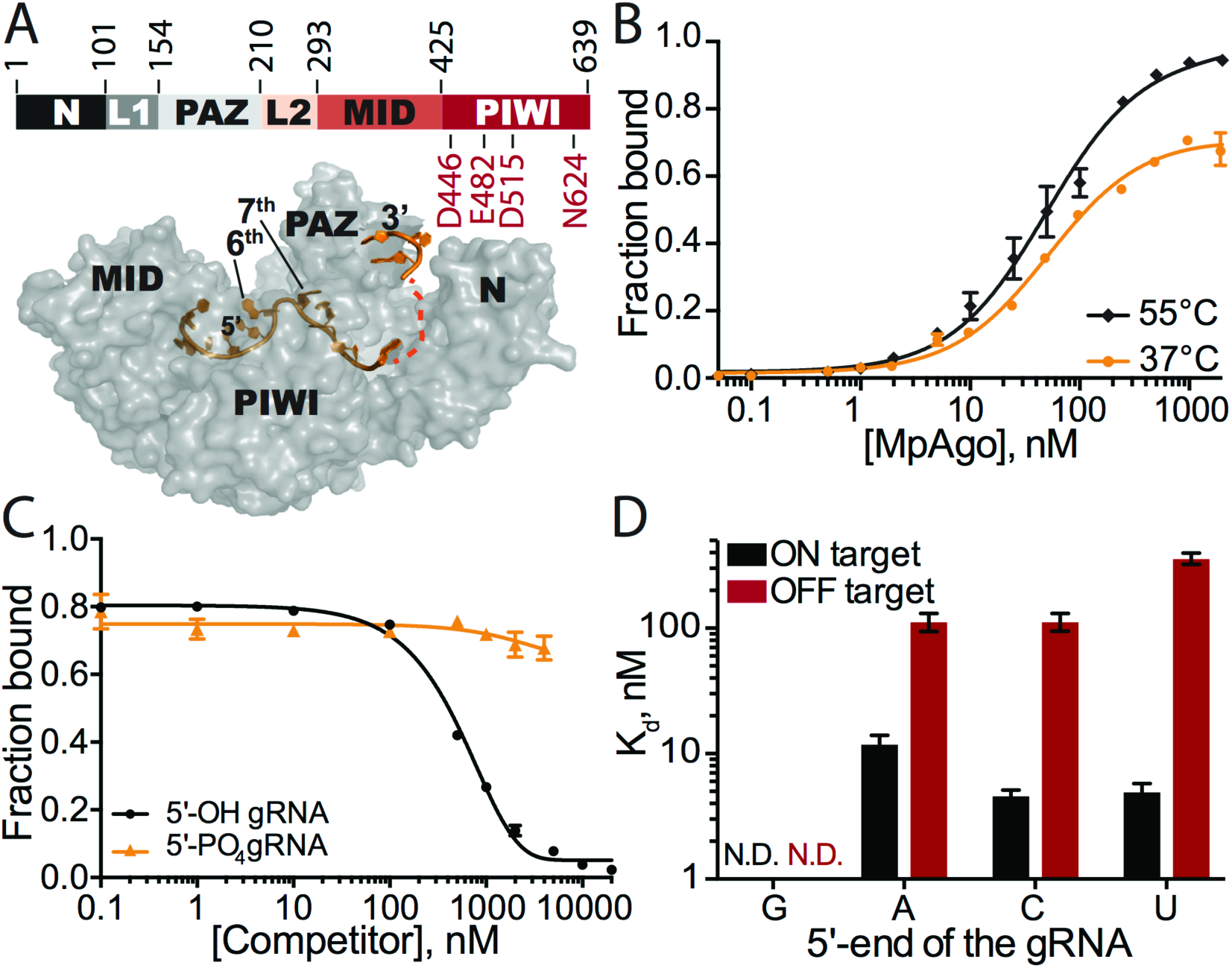
Stability and specificity of *in vitro* reconstituted MpAgo RNPs. **(A)** (Top) Organization of MpAgo’s domains and (bottom) the structure (PDB ID: 5I4A) of MpAgo (grey) bound to its guide RNA (gRNA, orange). The bi-lobed structure includes a PAZ (PIWI-Argonaute-Zwille) domain that anchors the 3′-end of the gRNA, while the second lobe includes a MID (middle) and a PIWI (P-element Induced WImpy testis) domain that binds the 5’-end of the gRNA, and includes the catalytic residues, as marked. The gRNA bound to MpAgo is bent at the 6^th^ and 7^th^ nts. **(B)** MpAgo binding of gRNA at two temperatures: 37 °C (orange) and 55 °C (black). The fraction of bound gRNA is plotted as a function of MpAgo concentration. The data fit with a standard binding isotherm (solid lines) yield K_d_ values at 37 °C and 55 °C of 52 ± 4 nM and 49 ± 5 nM, respectively. Data are represented as the mean ± standard deviation (SD) from three independent experiments. **(C)** Competition-binding assay at 37 °C to examine displacement of gRNA from the MpAgo RNP complex, with 5’-OH RNA competitor (black) and 5’-PO_4_ RNA competitor (orange). The adjusted half-inhibitory concentrations (IC_50_) are 0.25 ± 0.08 μM and > 4.5 μM for the 5’-OH RNA and 5’-PO_4_ RNA competitors, respectively. Data are represented as the mean ± SD from three independent experiments. **(D)** Substrate ssRNA filter-binding assays with catalytically inactive MpAgo RNP formed with a 5’-G, 5’-A, 5’-U, or 5’-C gRNA and either fully complementary substrate (ON target, black) or non-complementary ssRNA substrate (OFF target, red) in the presence of 2 μg/mL of heparin. The obtained average K_d_ values are plotted as the mean ± SD from three independent experiments. N.D. stands for ‘Not Detected.’

Here we present biochemical analyses of guide-dependent RNA targeting by MpAgo. We found that MpAgo is easily programmed with gRNAs to form RNA-protein complexes (RNPs). Reconstituted MpAgo RNPs bound fully complementary RNA substrates with moderate affinity and specificity. However, incorporation of 5-bromo-2′-deoxyuridine (BrdU) at the 5’ end of gRNAs stabilized the MpAgo-gRNA complex (MpAgo RNP), and greatly increased MpAgo RNP affinity and specificity in binding substrate RNAs. Moreover, MpAgo RNPs with 5’-BrdU gRNAs selectively bound substrates that differ by only one nucleotide, which enabled isolation of inosine-edited RNA substrates from a complex mixture.

## RESULTS

### Reconstituted MpAgo-gRNA complexes bind fully complementary RNA substrates with moderate affinity and specificity

The CRISPR locus associated MpAgo can be programmed with a short 5’-OH gRNA to cleave single stranded DNA and RNA substrates *in vitro* (16). We used catalytically active MpAgo for cleavage assays and catalytically inactive MpAgo for binding assays (16). Since the endogenous guides and targets for *M. piezophila* are not known, we investigated MpAgo interactions with a previously published 5’-OH 21-nucleotide (nt) gRNA (***SI Appendix***, Table S1) (16). We determined the equilibrium dissociation constant of MpAgo for its gRNA at 55 °C using a filter binding (FB) assay, since *M. piezophila* grows at 45 - 70 °C (17). We observed that MpAgo binds the 5’-OH gRNA with ~50 nM affinity (Fig. 1B). MpAgo RNP reconstituted at 37 °C, which would be compatible with expression and assembly of MpAgo RNPs in mesophilic (i.e. human) cells, bound the 5’-OH gRNA at 37 °C with a similar affinity as that at 55 °C, although with slightly diminished overall binding (Fig. 1B).

To assess the stability of the reconstituted MpAgo RNP *in vitro*, preassembled MpAgo RNP complexes were titrated with either excess 5’-OH or 5’-phosphorylated competitor gRNAs of identical sequence. We first measured the off-rate (k_off_) of 5’-OH gRNAs pre-bound to MpAgo at 55 °C to be approximately 0.03 min^−1^, with a small population of MpAgo RNPs remaining stable for long times in the presence of excess competitor gRNA (***SI Appendix***, Fig. S1A). Using incubations of about 45 times the half-life of the MpAgo-gRNA complex, the observed half-inhibitory concentrations (IC_50_) for 5’-OH and 5’-phosphorylated gRNAs reveal that MpAgo has a very low affinity for 5’-phosphorylated gRNAs (Fig. 1C), in agreement with published data (16). The IC_50_ values also demonstrate that the 5’-OH gRNA is not easily displaced from MpAgo RNP assembled at 55 °C, by either 5’-hydroxylated or by 5’-phosphorylated RNAs. We obtained similar results for the stability and 5’-end specificity of MpAgo RNPs assembled at 37 °C (***SI Appendix***, Fig. S1A).

We then tested the stability of MpAgo RNPs reconstituted at 37 °C and 55 °C in the presence of RNA targets. MpAgo RNPs immobilized on beads through a biotin tag on the 3’ end of a 33-nt gRNA were incubated with a 10-fold excess of mRNA containing a fully complementary target site (ON-target sample), or an excess of total RNA purified from HEK293T cells (OFF-target sample) (***SI Appendix***, Fig. S2A). Notably, in the presence of a fully complementary RNA target, we observed more than 50% displacement of MpAgo from MpAgo RNPs assembled at 37 °C, and nearly 30% displacement from MpAgo RNPs reconstituted at 55 °C (***SI Appendix***, Fig. S2B). However, using 22-nt gRNAs (radiolabeled on the 3’ end), MpAgo RNPs were stable in the presence of ON-and OFF-target RNAs (***SI Appendix***, Fig. S2C). Collectively, these results indicate that the temperature of the reconstitution reaction does not affect the stability of the MpAgo RNPs. However, binding of fully complementary target RNAs to MpAgo RNPs with gRNAs that have longer 3’ extensions can destabilize the RNPs (***SI Appendix***, Figs. S2B, D).

Previously, it was shown that MpAgo does not have a preference for the 5’-terminal nucleotide of the gRNA in targeting ssDNA substrates for cleavage (16). Since the cellular target of MpAgo is not known, and some Argonautes including MpAgo target ssRNA *in vitro* (16), we tested whether the 5’-end nucleotide of the gRNA affects the ability of MpAgo RNPs to bind and cleave RNA. Regardless of the gRNA 5’ nucleotide identity (***SI Appendix***, Fig. S3A), MpAgo cleaved ssRNA substrates with similar rates and efficiencies (***SI Appendix***, Fig. S3B). We then examined if the 5’-terminal nucleotide of the gRNA influences ssRNA substrate binding efficiency and specificity *in vitro*, using an RNA substrate that was either fully complementary (ON target) or non-complementary (OFF target) to the gRNA, over a range of MpAgo RNP concentrations (Fig. 1D; ***SI Appendix***, Figs. S3C, F). Whereas ON target binding efficiencies of RNPs with 5’-A, C or U gRNAs are comparable, the specificity is affected by the identity of the 5’-terminal position of the gRNA. RNPs programmed with 5’-U gRNAs are more specific than the RNPs programmed with 5’-A and 5’-C gRNAs, whereas MpAgo RNPs programmed with a 5’-G gRNA bind fully complementary target RNAs inefficiently. The 5’-nt preference of MpAgo for 5’-U gRNAs is not due to differences in gRNA binding affinity, as we found the MpAgo affinities for gRNAs to be similar with all four 5’-nucleotides (***SI Appendix***, Fig. S3G). The fact that MpAgo RNP cleaves only complementary substrates (16) indicates that non-specific binding of the ssRNA substrates obtained here is likely gRNA independent and could be caused by incomplete assembly of the MpAgo RNP, allowing non-specific RNA binding to the gRNA binding cleft of free MpAgo (16).

### Chemical modification of the 5’-terminal nucleotide of the gRNA improves MpAgo RNP specificity and affinity for ssRNA substrates

Despite low nanomolar affinity of the MpAgo RNP for ssRNA substrates, the moderate specificity of MpAgo RNPs (Fig. 1D) prompted us to engineer a more specific MpAgo RNP. The previously published structure of MpAgo revealed that the 5’-terminal nucleotide of the bound gRNA is flipped into a pocket formed by MpAgo’s MID and PIWI domains (Fig. 1A), where it stacks on the aromatic ring of tyrosine 379 (Y379) (Fig. 2A) (16). We tested whether the close proximity of the 5’-terminal nucleotide and the Y379 aromatic ring would favor crosslinking of the gRNA to MpAgo. We used a gRNA containing 5-bromo-2′-deoxyuridine (BrdU) at the 5’-end position (***SI Appendix***, Table S1) and crosslinked it to the protein using 305 nm UV light.

**Figure 2.**
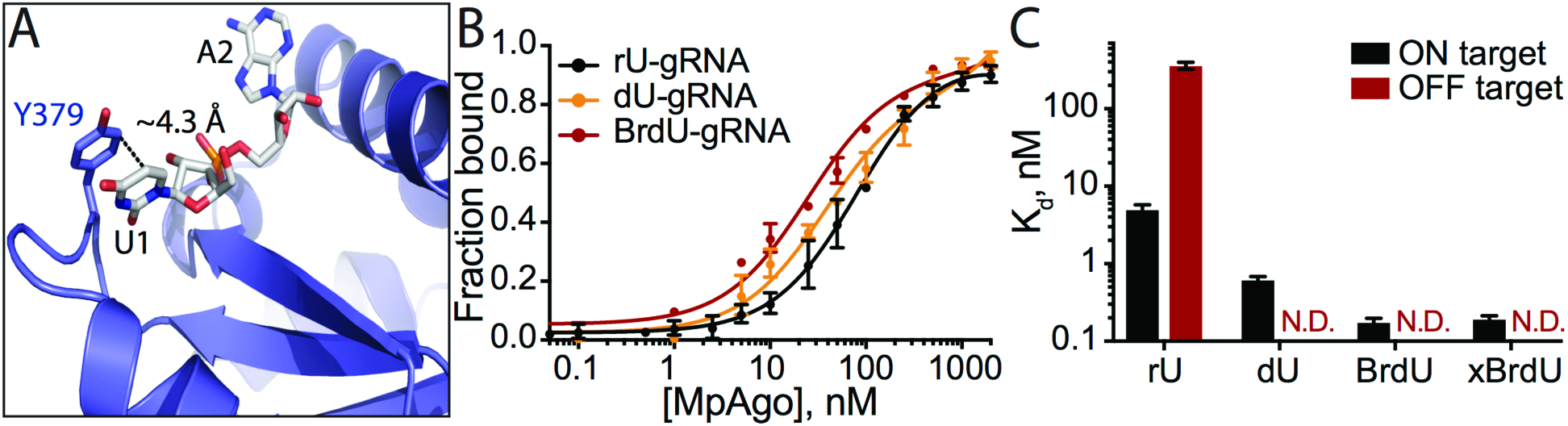
MpAgo RNP assembly using 5’-end modified gRNAs and specificity for binding complementary ssRNA substrates. **(A)** MpAgo (blue) (PDB ID: 5I4A) forms a binding pocket for the 5’-end nucleotide (U1) of the gRNA (grey) that stacks on the aromatic ring of tyrosine residue Y379. **(B)** MpAgo and gRNA binding curves obtained for 5’-end ribonucleotide (rU, black), 2′-deoxyuridine (dU, orange), and 5-bromo-2′-deoxyuridine (BrdU, red) containing gRNAs at 55 °C. The fraction of bound gRNA is plotted as a function of MpAgo concentration. The data fit with a standard binding isotherm (solid lines) reveal MpAgo binds the rU, dU and BrdU gRNAs at 55 °C with average K_d_ values of 81 ± 9 nM, 37 ± 6 nM and 25 ± 3 nM, respectively. Data are represented as the mean ± standard deviation (SD) from three independent experiments. **(C)** The average K_d_ values of the MpAgo RNPs for either fully complementary substrate (ON target, black) or non-complementary ssRNA substrate (OFF target, red) were obtained using MpAgo RNPs formed with a rU, dU, or BrdU gRNA, or with a BrdU gRNA with subsequent 1h exposure to 305 nm UV light for gRNA crosslinking to MpAgo (xBrdU). The obtained K_d_ values are plotted as the mean ± SD from three independent experiments. N.D. stands for ‘Not Detected.’

Although UV crosslinking reached ~22%-33% efficiency (***SI Appendix***, Fig. S4), analysis of non-crosslinked MpAgo RNPs revealed that the 5’-BrdU modification alone was sufficient to improve MpAgo RNP affinity and specificity for substrate RNAs (Fig. 2C; ***SI Appendix***, Fig. S5). Replacing the 5’-end ribonucleotide (rU) of the gRNA with 2′-deoxyuridine (dU) or BrdU slightly increased MpAgo affinity for the gRNA, with each modification making a positive contribution (Fig. 2B). To assess the effect of these 5’-end gRNA modifications on MpAgo RNP stability, we measured the off-rates (k_off_) of 5’-end dU and BrdU gRNAs pre-bound to MpAgo at 55 °C to be approximately 0.024 min^−1^ and 0.011 min^−1^, with a small population of MpAgo RNPs remaining stable for long times in the presence of excess competitor gRNA (***SI Appendix***, Fig. S5A). These data indicate that the BrdU modification stabilizes the MpAgo RNP. Interestingly, in optimized conditions (1:1.5 MpAgo:gRNA ratio, and 2 μg/mL of heparin), a dU at the gRNA 5’-end increased MpAgo RNP affinity for a fully-complementary ssRNA substrate (21-nt complementarity) by 10 times (***SI Appendix***, Figs. S3F, S5B), and a 5’-BrdU further increased MpAgo RNP affinity for the ssRNA substrate to better than 200 pM (***SI Appendix***, Fig. S5C). Notably, these substitutions also decreased OFF-target binding to undetectable levels (Fig. 2C; ***SI Appendix***, Figs. S5B-C). In the case of 5’-BrdU, these improvements in target binding did not depend on UV crosslinking (Fig. 2C; ***SI Appendix***, Figs. S5C-D). The following experiments were carried out with UV-crosslinking, however the above results indicate this is unnecessary for 5’-BrdU gRNAs to exert their effects on MpAgo RNPs.

### MpAgo utilizes a seed region on the gRNA with highest sensitivity at the 3^rd^, 6^th^, 7^th^ and 9^th^ nucleotides

The seed sequence in the gRNA contributes greatly to the specificity of Ago RNP interactions with target substrates. The canonical seed region of all eAgos and some pAgos is composed of the 2^nd^ to 8^th^ nts of the gRNA and has an A-form helical conformation in the Ago RNP complex (18–23). Furthermore, the 2^nd^ to 4^th^ bases of the seed sequence are exposed to the solvent for interaction with the complementary substrate (22). Notably, in contrast to eAgos and pAgos, the previously determined structure of an MpAgo-gRNA complex revealed that the gRNA and the predicted seed region have a unique conformation, a sharp kink between the 6^th^ and 7^th^ nucleotide of the gRNA that disrupts continuous A-like helical base stacking of the 2^nd^ – 8^th^ nucleotides (16). Interestingly, the tolerance for dinucleotide mismatches across the MpAgo gRNA when targeting ssDNA revealed that the mismatches at the 5^th^ to 15^th^ nts of the gRNA abolish the cleavage of ssDNA substrates the most (24). These structural and biochemical studies suggest that the seed region of the MpAgo may differ from the canonical seed used by previously characterized eAgos and pAgos.

We investigated whether mismatches of 3 or 6 nucleotides between ssRNA target and nucleotides at the 3′-end of 5’-BrdU gRNAs affect MpAgo RNP binding to ssRNA substrate (Fig. 3A). Surprisingly, in optimized conditions (1:1.5 MpAgo:gRNA ratio, and 10 μg/mL of heparin (***SI Appendix***, Figs. S6A, B) reducing complementarity between the ssRNA and gRNA to 18 nts or 15 nts enhanced the affinity of MpAgo RNP substantially, with K_d_ values of ~100 pM (Fig. 3B). Due to the high affinity of 15-nt target RNAs (Fig. 3B), we mapped the seed region of MpAgo when targeting ssRNA substrates using a baseline of 15-nt complementarity to target ssRNAs (Fig. 3A). We introduced single nucleotide mismatches in the RNA substrate at the positions corresponding to the 1^st^ to 15^th^ nucleotides in a 21 nucleotide long gRNA, keeping the sequence of the gRNA constant (***SI Appendix***, Table S1, Fig. S7A) and determined the equilibrium dissociation constants using filter binding assays (Fig. 4A; ***SI Appendix***, Fig. S7B). Due to the location of the 1^st^ gRNA nucleotide in a separate binding pocket (Fig. 2A), there is no effect of a mismatch at the first position. We noticed that the K_d_ values for the substrates that have mismatches spanning the 2^nd^ to 12^th^ positions (with the exception of the 9^th^ position), are at least 5 times higher relative to the K_d_ value for the fully complementary ssRNA substrate (Fig. 4A).

**Figure 3.**
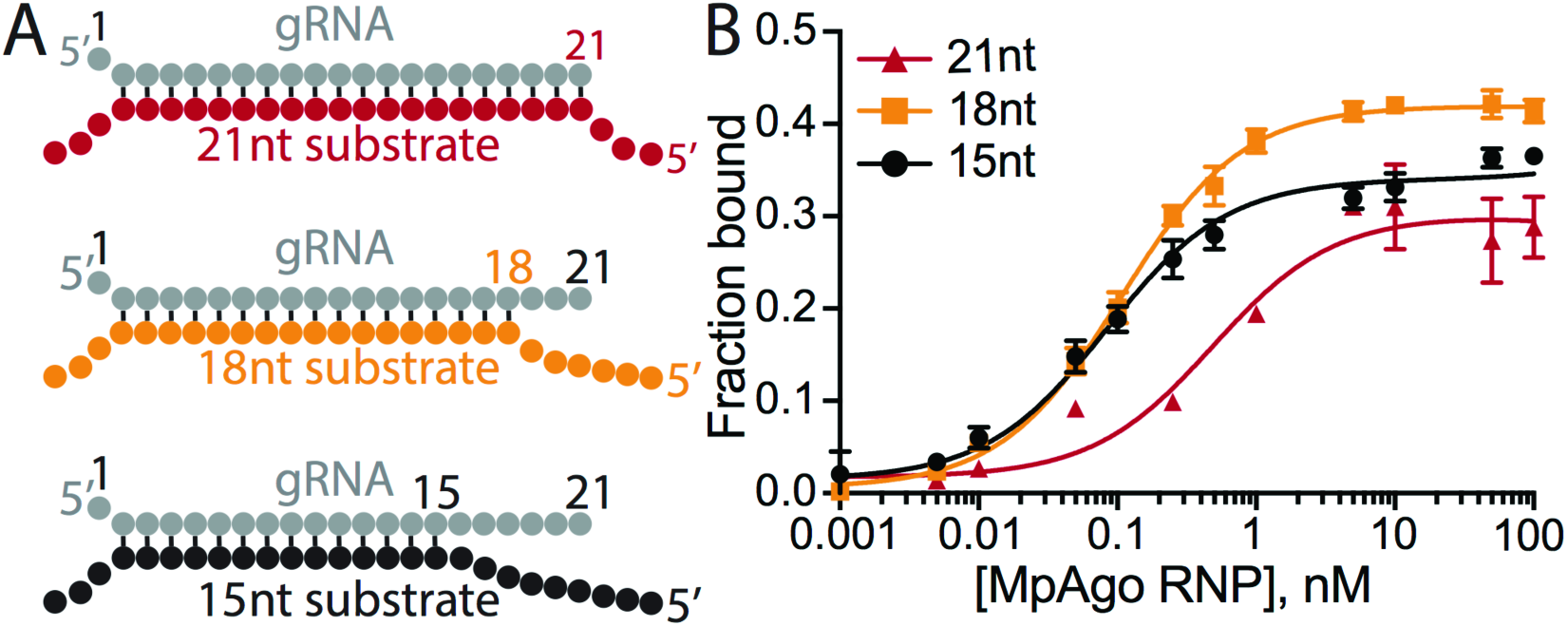
MpAgo RNP programmed with 5’-BrdU gRNA binds ssRNA substrates with high affinity. **(A)** Cartoon of three duplexes of the gRNA and different ssRNA substrates with varying degrees of complementarity to the gRNA used in binding assays: full 21-nt complementarity (top, red), 18-nt complementarity with 3 mismatches at the 3′-end of the gRNA (middle, orange), and 15-nt complementarity with 6 mismatches at the 3′-end of the gRNA (bottom, black). The first nt at the 5’-end of the gRNA does not base pair due to MpAgo interactions (Fig. 2A). **(B)** Filter-binding assays with 21-nt complementarity (red), 18-nt complementarity (orange), or 15-nt complementarity (black) in the presence of 10 μg/mL of heparin. The amount of bound ssRNA substrate is plotted as a function of MpAgo RNP concentration. The obtained average K_d_’s for MpAgo RNP and the substrates with 21-, 18-, and 15-nt complementarity are 500 ± 115 pM, 107 ± 6 pM, and 83 ± 8 pM, respectively. Data are represented as the mean ± SD from three independent experiments.

**Figure 4.**
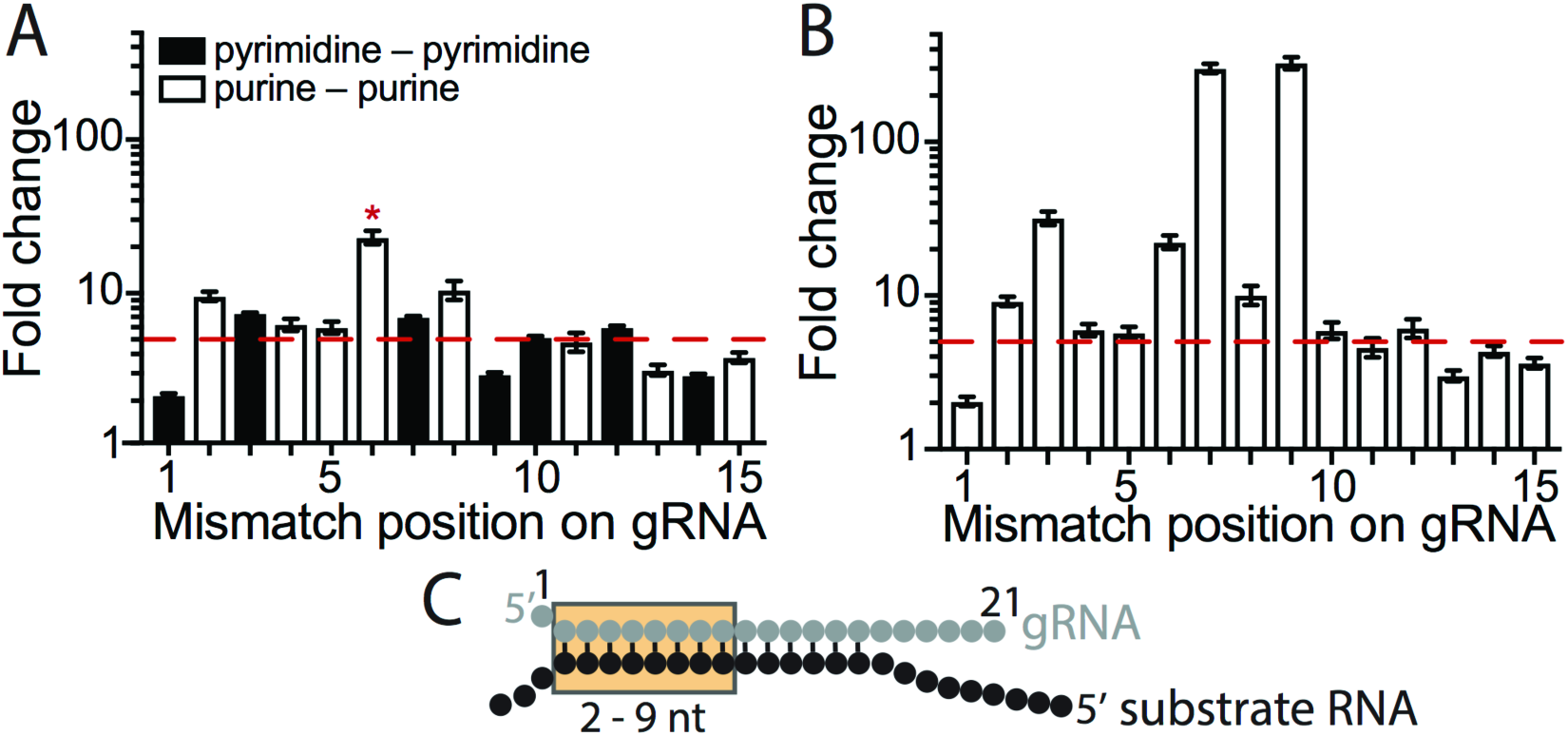
Determination of the gRNA seed region of MpAgo RNPs. **(A)** Substrate ssRNA filter-binding assays with ssRNA substrates that contain a single nucleotide mismatch along the gRNA from the 1^st^ to 15^th^ position. The sequence of the gRNA was kept constant, and the ratios of the average K_d_’s of the substrates that create mismatch (K_d_^mm^) to the fully complementary substrate (K_d_^Compl^) (Fold change = K_d_^mm^/ K_d_^Compl^) were plotted against the position of the mismatch. The red dash line indicates the value equal to ~5 times the K_d_ for the fully complementary ssRNA substrate. Asterisk (*), A-G mismatch at the 6^th^ position (guide RNA-substrate RNA). The K_d_ for a G-A mismatch at the 6^th^ position is 779 ± 71 pM, using a different gRNA (Table S1). **(B)** The ratios of the average K_d_’s of the substrates that create purine-purine mismatch (K_d_^mm^) to the fully complementary substrate (K_d_^Compl^) (Fold change = K_d_^mm^/ K_d_^Compl^) were plotted against the position of the mismatch. The red dash line indicates a value equal to 5. The purine-purine mismatches at the 2^nd^ through 12^th^ positions decrease the affinity of MpAgo RNP more than 5 fold. The 3^rd^, 6^th^, 7^th^ and 9^th^ positions reduce the affinity of MpAgo RNP the most, 32-, 22-, 301- and 327-fold respectively. **(C)** The gRNA-substrate RNA duplex with the seed region (orange box).

The type of mismatch also affects the observed K_d_ value for target binding. In the context of scanning target mismatches to one gRNA, purine-purine mismatches (A-G mismatch in gRNA-substrate at the 6^th^ position, and G-A mismatches at the 2^nd^ and 8^th^ positions) in general weakened target binding substantially more than pyrimidine-pyrimidine mismatches (Fig. 4A; ***SI Appendix***, Fig. S5). To check whether the negative impact of the mismatch at the 6^th^ position is due to the nature of an A-G mismatch, we exchanged the gRNA sequence to obtain a G-A mismatch at the 6^th^ position, which resulted in a K_d_ very similar to those for substrates that form G-A mismatches at the 2^nd^ and 8^th^ positions (Fig. 4A; ***SI Appendix***, Fig. S9D). We used additional gRNAs and substrates (***SI Appendix***, Table S1) to probe purine-purine mismatches at the 3^rd^, 7^th^, 9^th^, 10^th^, 12^th^ and 14^th^ positions to more rigorously define the seed region for MpAgo RNPs (***SI Appendix***, Fig. S8). The observed K_d_ values indicate that the MpAgo RNP seed region spans nucleotides 2 - 9, and is highly sensitive to mismatches at the 3^rd^, 6^th^, 7^th^ and 9^th^ positions (Fig. 4B). Taken together, our data suggests that MpAgo RNP has an extended seed compared to the canonical seed in eAgos and some pAgos (18–23), when targeting ssRNAs (Fig. 4C).

### MpAgo RNPs can be programmed to distinguish between single nucleotide variants

Using the 6^th^ and 7^th^ nucleotides located at the kink of the gRNA (16) (Fig. 1A), we investigated whether MpAgo RNPs could distinguish between RNA substrates that differ by only one nucleotide at the corresponding complementary position. Using MpAgo RNPs programmed with 5’-BrdU gRNAs that have A, G, U or C either at the 6^th^ or at the 7^th^ position and RNA substrates with all permutations at the corresponding positions (***SI Appendix***, Figs. S9, S10), we measured average dissociation constants for the 6^th^ and 7^th^ position (Figs. 5A, B). Notably, at the 6^th^ position MpAgo RNP has the highest affinity for the fully complementary substrates (gRNA:substrate base pairs G:C, U:A, A:U and C:G), substrates which form pyrimidine-pyrimidine mismatches (U–U and C–U), and the substrate that forms a G•U wobble. High-affinity binding of substrates with U–U and C–U mismatches suggesting that these mismatches are well accommodated in the gRNA-substrate RNA duplex at this position. Interestingly, only the G:C base pair forming substrate binds more strongly than the substrate that creates a G•U wobble, whereas the binding affinities of the substrates that form U:A, A:U and C:G Watson-Crick base pairs are ≥ 2 times weaker. Purine-purine mismatches destabilize substrate RNA binding most, with A–A, A–G, G–A and G–G mismatches increasing the K_d_ values 20- to 56fold relative to Watson-Crick pairs. Moreover, MpAgo RNP binding affinity to ssRNA also depends on which nucleotide of the mismatch is on the gRNA strand. The binding affinity of the substrate forming a G•U wobble is 21 times tighter than that of the substrate forming a U•G wobble. Similar trends are observed for other types of mismatches, e.g. G–A vs. A–G, C–A vs. A–C, and C–U vs. U–C (Fig. 5A). Even in the case of perfectly matching gRNA-substrate duplexes, a C:G base pair binds 13 times more tightly than a G:C base pair.

**Figure 5.**
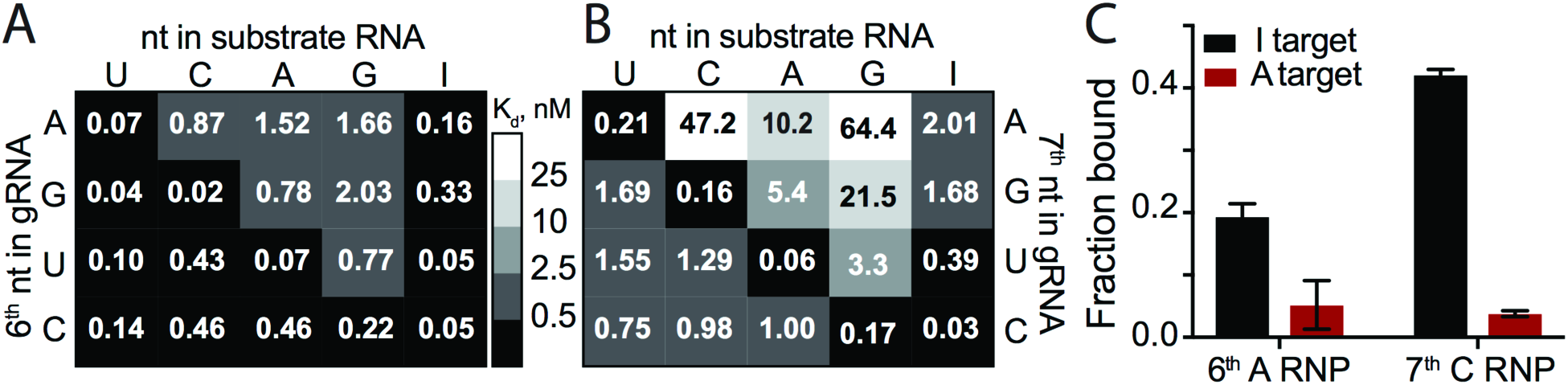
5’-BrdU-gRNA MpAgo RNP ssRNA substrate binding specificity at the 6^th^ and 7^th^ positions. Filter-binding assays were performed using 5’-BrdU-gRNA MpAgo RNPs programmed with A, G, U or C gRNA at the 6^th^ **(A)** or at the 7^th^ **(B)** positions and ssRNA substrates containing U, C, A, G or I at the corresponding position. The average K_d_’s were extracted and are represented in the heat maps. **(C)** A filter-binding assay to isolate the A-to-I edited substrate (black) but not the non-edited substrate (red) from 500 ng of total RNA purified from HEK239T cells, in the presence of 200 ng/μL of yeast tRNA. For this assay MpAgo RNP programmed with a 5’-BrdU gRNA containing either A at its 6^th^ or C at its 7^th^ position was used. The fraction of the ssRNA substrate bound to MpAgo RNP was quantified and plotted as the mean ± SD from three independent experiments.

In contrast to the 6^th^ position, at the 7^th^ position MpAgo RNP has the highest affinity only for complete complementary substrates across 15 nucleotides (gRNA:substrate base pairs G:C, U:A, A:U and C:G). Even though pyrimidine-pyrimidine mismatches and the G•U and U•G wobble affect the binding affinities the least, they increase the K_d_ values ~10- to 20-fold relative to Watson-Crick pairs, suggesting that even these mismatches are not well accommodated in the gRNA-substrate RNA duplex at this position. Purine-purine mismatches (A–A, A–G, G–A and G–G) and purine-pyrimidine mismatch A–C destabilize substrate RNA binding even more, increasing the K_d_ values 10- to 100-fold relative to Watson-Crick pairs. Similarly to the 6^th^ position, MpAgo RNP binding affinity to ssRNA also depends on which nucleotide of the mismatch is on the gRNA strand. The binding affinity of the substrate forming a C–A mismatch is 47 times tighter than that of the substrate forming an A–C mismatch. Similar trends are observed for other types of mismatches, e.g. G–A vs. A–G, G•U vs. U•G, and C–U vs. U–C (Fig. 5B). Collectively, our analysis of the 6^th^ and 7^th^ positions of the gRNA and corresponding positions of the ssRNA substrate indicate that MpAgo monitors not only the type of mismatch, but also the identity of the bases forming the mismatch.

Next we tested whether MpAgo RNPs have high enough specificity to recognize modified nucleotides in RNA substrates. For instance, adenosine deaminases acting on RNA (ADARs) catalyze adenosine to inosine (A–to–I) editing of certain mRNAs (25, 26). Compared to adenosine, inosine has very different base-paring properties (27). We tested whether MpAgo RNPs could be programmed to distinguish between A-to-I edited and non-edited RNA substrates using MpAgo RNPs programmed with gRNAs that have A, G, U or C at either the 6^th^ or at the 7^th^ positions and substrate with inosine at the corresponding complementary positions (Figs. 5A, B; ***SI Appendix***, Figs. S9, S10). Although MpAgo RNPs have low picomolar affinity for the inosine containing substrate irrespective of which nucleotide is used at the 6^th^ position of the gRNA, the presence of an A at the 6^th^ position of the gRNA leads to ~10-fold weaker binding to substrates with an A instead of I (Fig. 5A; ***SI Appendix***, Fig. S9). Intriguingly at the 7^th^ position, MpAgo RNP binds the I-containing substrate weaker than at the 6^th^ position with the exception of the MpAgo RNP that has C at the 7^th^ position (Fig. 5B; ***SI Appendix***, Fig. S10). Moreover, the presence of a C at the 7^th^ position of the gRNA leads to ~33-fold weaker binding to substrates with an A instead of I (Fig. 5B; ***SI Appendix***, Fig. S10).

We then investigated whether MpAgo RNP programmed with the 5’-BrdU gRNA containing either A at its 6^th^ or C at its 7^th^ position could be used to isolate A-to-I edited RNAs from a complex mixture. We spiked in either a radiolabeled I-containing RNA substrate or a radiolabeled A-containing RNA substrate into HEK293T total RNA, and incubated the mixtures with MpAgo RNP programmed with a gRNA containing either A at its 6^th^ or C at its 7^th^ position. We also substituted bulk tRNA for heparin in the binding reactions to make the conditions of binding reactions more physiological. At the 6^th^ position, MpAgo RNP bound an I-containing substrate with ~19% efficiency, while the A-harboring RNA substrate was bound only with 5% efficiency. At the 7^th^ position, MpAgo RNP bound an I-containing substrate with ~42% efficiency, while the A-harboring RNA substrate was bound only with 4% efficiency (Fig. 5C). These results are consistent with the specificity seen with pure substrates in FB assays (***SI Appendix***, Figs. S9, S10) at the concentration of MpAgo RNP used with the complex mixture (500 pM MpAgo RNP). These results demonstrate that MpAgo RNPs can distinguish between the RNA substrates that differ by only one nucleotide with high specificity, and that it can be programmed to isolate A-to-I edited substrates from complex mixtures.

## DISCUSSION

As opposed to all other known Agos, the CRISPR-associated Agos in the *M. piezophila* family bind chemically distinct 5’-hydroxylated gRNAs rather than 5’-phosphorylated guides. This implies that the origin and biological assembly pathway of these Ago RNPs are also likely to be unique. It is possible that MpAgo gRNAs are generated by nucleases associated with the CRISPR operon, namely Cas6, Csx1, or Csm6 (28–30). All three nucleases produce 5’-hydroxylated short RNAs (31–33) that could serve as MpAgo guides. We speculate that either Csx1 or Csm6 but not Cas6 is responsible for generating gRNAs for MpAgo, as Cas6 produces short RNAs that contain a conserved (8-nt) repeat sequence derived from the CRISPR array at their 5’ end (32). The conserved sequence spans the most important region of the seed sequence (Fig. 4), which would defeat the functional role of this region of the gRNA. The fact that the source of guides is likely unique for these CRISPR-associated Agos suggests that their biological role could differ from other pAgos (8, 13–15), a question that will require future experiments to answer.

In addition to the unknown origin of MpAgo gRNAs, the physiological targets for MpAgo are currently unclear. We showed that MpAgo cleaves ssRNA substrates with a cleavage rate of 0.39 - 0.64 min^−1^, similar to rates observed for ssDNA substrate cleavage (k_obs_ ~0.55 min^−1^ (16)). This implies that both ssRNA and ssDNA may be physiological targets of MpAgo. We also found that MpAgo, once formed as an RNP, has moderate affinity (~5 nM K_d_) and specificity for target RNAs (Fig. 1D; ***SI Appendix***, Figs. S3D-F). Remarkably, simple substitution of the 5’-end nucleotide of the gRNA with 5’-dU greatly increases the affinity and specificity of MpAgo RNPs for RNA substrates (***SI Appendix***, Figs. 2C, S5B). Although intended for UV crosslinking, use 5’-BrdU gRNAs without crosslinking further increases MpAgo RNPs affinity and specificity for target RNAs (Fig. 2C; ***SI Appendix***, Fig. S5C). Moreover, MpAgo RNPs programmed with 5’-BrdU gRNAs are twice as stable compared to the RNPs programmed with 5’-rU gRNAs (***SI Appendix***, Figs. S1, S5A) and can bind complementary RNA substrates over 300 times more efficiently than non-complementary substrates with a mismatch at a single nucleotide (Fig. 4B).

The specificity of the Ago RNP’s interaction with its substrates is determined by the seed sequence in the guide RNA. We have shown the MpAgo RNP seed region spans the 2^nd^ to 9^th^ nucleotides of the gRNA when targeting ssRNA. This is distinct from the extended seed region when MpAgo targets ssDNA substrates (24), but similar to the seed region reported for other Agos, with the 3^rd^, 6^th^, 7^th^ and 9^th^ positions affecting MpAgo RNP’s specificity in binding target RNAs the most. With eAgos, it has been reported that the 4^th^ and 5^th^ nt of the gRNA contribute the most to specificity (34), although other reports suggest central positions of the guide RNA can also confer specificity (35). In MpAgo, the 6^th^-7^th^ positions, may serve as conformational checkpoints, since the guide RNA is kinked between the 6^th^ and 7^th^ nucleotide (Fig. 1A). We hypothesize that base pairing of the 6^th^ and 7^th^ nucleotides releases the distortion in the gRNA and allows further 5’ to 3’ propagation of the guide-substrate base pairing. The structure of MpAgo RNP bound to a DNA substrate suggests that two protein helices of the L2 domain and a loop of the PIWI domain could enable MpAgo to sense the shape of the minor groove at the 6^th^ and 7^th^ position of the gRNA and ensure that the gRNA-RNA substrate duplex is complementary (***SI Appendix***, Fig. S11) (24), although MpAgo RNP interactions with DNA and RNA substrates are likely to be distinct (Fig. 4). Analysis of substrate specificity at the 6^th^ and 7^th^ nt of the gRNA demonstrate that MpAgo not only monitors the position and type of the mismatch but also the identity and position of the bases forming the mismatch. This discovery reveals a coding feature of the MpAgo RNP that enables it to discriminate between substrate RNAs differing by only a single nucleotide.

Interestingly, the MpAgo RNP can also be programmed to distinguish between target RNAs that are post-transcriptionally modified, i.e. that contain common base modifications such as inosine. Despite inosine’s ability to base pair with all four canonical bases, it forms the most stable base pair with cytidine. Therefore, it is recognized as guanosine by the translational machinery when it is present in coding mRNA sequences and can cause amino acid substitutions (36). Additionally, perturbations of A-to-I editing correlate with several neurological diseases (37). Despite the prevalence of A-to-I editing, the functions of most editing sites remain unknown. The method primarily used to identify editing sites is based on the comparison of cDNA and corresponding genomic DNA sequences (36). However, there is no reliable method to investigate the function, cellular localization, and interaction networks of specific, edited RNA transcripts *in vivo*. The ability of MpAgo RNPs to isolate A-to-I edited RNAs from a complex mixture should enable deeper exploration of the biological function of inosine modifications in RNAs. In principle, MpAgo RNPs could also be used to isolate mRNAs with C-to-U deaminations (38) using gRNAs with an A at the 7^th^ position (Fig. 5B).

MpAgo has moderate affinity for 5’-hydroxylated gRNAs (K_d_ ~50 - 100 nM for all RNA guides that contain ribonucleotide at the 5’-end, and ~25 nM for 5’-BrdU guides), when compared to other Agos (K_d_ ~1 − 7 nM for *Rhodobacter sphaeroides* Ago (39) and human Argonaute 2, respectively (40)). Notably, the higher affinity of MpAgo RNPs for ssRNA substrates (K_d_ ~5 nM for RNPs programmed with 5’-ribo gRNAs, and ~0.02 - 0.5 nM for RNPs programmed with 5’-BrdU guides) than of MpAgo for gRNAs (K_d_ 50 - 104 nM for 5’-ribo gRNAs, and ~25 nM for 5’-BrdU gRNAs) implies that binding of gRNA to MpAgo is stabilized by binding complementary RNA substrates (Figs. 1B, D; Figs. 2B, D; Figs. 5A, B). Thus, MpAgo is relatively easily programmed with defined gRNAs to target diverse RNA substrates of interest, and furthermore can be programmed at 37 °C, suitable for use in mesophilic cells. MpAgo RNPs with 5’-BrdU gRNAs bind complementary ssRNA substrates with much higher affinities than most RNA binding proteins currently employed for RNA targeting. Whereas the MS2 and PP7 coat proteins, the λN protein, repurposed RCas9, and more recently characterized type II-A Cas9 bind their RNA substrates with 1.6 nM – 5 nM affinities, only the CRISPR nuclease Csy4 binds its RNA hairpin substrate with similar affinity to MpAgo, ~50 pM (41–46). Of these systems, only *S. pyogenes* and *S. aureus* Cas9 are easily programmable, and *Streptococcus pyogenes* Cas9 requires a PAMmer DNA oligonucleotide for efficient RNA binding (44, 46–48). Notably, the cellular concentration of mRNAs in mammalian cells is generally in the 10 – 100 pM range, except for the most abundant transcripts (49, 50), matching the affinity range for MpAgo RNPs programmed with 5’-BrdU gRNAs. Finally, these MpAgo RNPs can be used to discriminate single nucleotide variants and to identify A-to-I edited target RNAs. In the future, it should be possible to use 5’-BrdU gRNA-loaded MpAgo RNPs to analyze, image and manipulate untagged RNAs in live cells with single nucleotide resolution.

## MATERIALS AND METHODS

A full description of the materials and methods used in this study, including *in vitro* RNA synthesis and radiolabeling, *in vitro* reconstitution of MpAgo RNP, MpAgo unloading assays, *in vitro* binding and cleavage assays, MpAgo-gRNA crosslinking experiments, A-to-I edited substrate isolation assays, and a list of oligonucleotides, is provided in the ***SI Appendix***.

## ACKNOWLEDGMENTS

We thank the members of the Cate and the Doudna laboratories, especially Steven C. Strutt, Emine Kaya, and Kevin W. Doxzen for their technical assistance and helpful discussions. A.L. acknowledges support from the Human Frontiers in Science Program (HFSP). This work was supported by the US National Institutes of Health (NIH) grant R01-GM065050 to J.H.D.C. and by a Frontiers Science award from the Paul Allen Institute to J.A.D. J.A.D. is an Investigator of the Howard Hughes Medical Institute (HHMI). The authors have submitted a patent related to this work.

## SUPPORTING INFORMATION

### SI MATERIALS AND METHODS

#### Protein Expression and Purification

Recombinant wild-type (WT) MpAgo (UniProtKB accession code H2J4R4) and catalytically inactive mutant D446A MpAgo (dMpAgo) were expressed in *E. coli* strain BL21 (DE3) (New England Biolabs) and purified as described previously (1). The purified proteins were concentrated to ~8 mg/mL, flash-frozen in liquid nitrogen, and stored at −80 °C. Catalytically active MpAgo was used for cleavage assays, whereas the catalytically inactive mutant was used in all binding assays. The abbreviation “MpAgo” is used for both versions of the protein.

#### RNA Synthesis and Labeling

RNAs longer than 29 nt were transcribed *in vitro* using synthetic DNA templates carrying a T7 promoter sequence (2). After transcription, the RNA was purified by denaturing 12% polyacrylamide gel electrophoresis (PAGE) and extracted from the gel using the crush-and-soak method followed by ethanol precipitation. The resulting transcribed RNAs were dephosphorylated using alkaline phosphatase CIP (NEB). After the reaction, the CIP was removed by phenol/chloroform extraction and the RNA was purified by ethanol precipitation. The RNAs were resuspended in RNase-free water and stored at −80 °C. Short and modified RNA and all DNA oligonucleotides were purchased from Integrated DNA Technologies (IDT). All nucleic acids used in this study are listed in **Table S1**.

For *in vitro* cleavage and binding assays, 10 pmol of RNA oligonucleotide was 5’-radiolabeled using [γ-^32^P] ATP (Perkin-Elmer) and 5 units of T4 polynucleotide kinase (NEB) in 1× T4 PNK buffer (NEB) at 37 °C for 30 min. The kinase was inactivated by incubation at 65 °C for 20 min and the labeling reaction was spun through an illustra^TM^ MicroSpin G-25 column (GE Life Sciences) to remove free nucleotides.

For the *in vitro* gRNA binding assay and the competition assays, 10 pmol of gRNA was 3′-radiolabeled using [^32^P] pCp (Perkin-Elmer) and 10 units of T4 RNA Ligase I (NEB). The labeling was performed overnight at 16 °C. The ligase was inactivated by incubation at 95 °C for 2 min and the reaction was purified from free nucleotides by spinning through an illustra^TM^ MicroSpin G-25 column (GE Life Sciences).

#### Assembly of the MpAgo RNP Complex

The MpAgo-gRNA complex was prepared by mixing MpAgo and gRNA with a ratio of 1:1.5 in RNP assembly buffer (20 mM Tris-HCl, 150 mM KCl, 1 mM MnCl_2_, 10 μg/mL BSA, 5% (v/v) glycerol, 2 mM DTT, pH_RT_ 7.5). RNP assembly buffer included 2 μg/mL heparin in Figs. 1–2, S1-S4. RNP assembly buffer included 10 μg/mL heparin in Figs. 3–5, S6–S10. After mixing the components, the samples were incubated either at 37 °C for 45 min or at 55 °C for 15 min and cooled down slowly to room temperature.

#### MpAgo-gRNA Filter-Binding Assays

The filter-binding assays were performed as described previously to obtain equilibrium binding isotherms (3). The gRNA binding experiments were performed in RNP assembly buffer described above. 0.1 nM of 3′-radiolabeled gRNA was incubated with increasing concentrations of WT MpAgo in a 50 μL reaction at either 37 °C for 45 min or at 55 °C for 15 min. 45 μL of each sample were applied to a dot-blot apparatus with low vacuum and three stacked membranes (Tuffryn (Pall Corporation), Protran (GE Healthcare), Hybond-N^+^ (GE Healthcare)) to separate MpAgo-gRNA complexes from free gRNA. Membranes were washed with washing buffer (20 mM Tris-HCl, 100 mM KCl, 5 mM MgCl_2_, pH_RT_ 7.5), air-dried and visualized by phosphor-imaging. The intensities of the free and MpAgo-bound gRNAs were analyzed using GelEval 1.35 (FrogDance Software). The obtained data of three independent experiments was plotted as function of MpAgo concentration and fit with a standard binding isotherm using the Prism 6 program (GraphPad Software, Inc.).

#### MpAgo Unloading Assay with streptavidin beads

The MpAgo RNP was prepared by mixing catalytically inactive MpAgo and biotinylated gRNA at a ratio of 1:2 in RNP assembly buffer. The samples were incubated either at 37 °C for 45 min or at 55 °C for 15 min and cooled down slowly to room temperature. 30 pmol of reconstituted RNP were combined with 60 μL of magnetic beads (Dynabeads MyOne Streptavidin M280; Life Technologies) in bead-binding buffer (10 mM Tris-HCl, 150 mM KCl, 2 mM MgCl_2_, 5% (v/v) glycerol, 0.05% (v/v) Tween 20, pH_RT_ 7.5). The binding reaction was incubated at 25 °C for 30 min on a rotating mixer. The beads were placed on a magnetic rack to remove the supernatant and were washed 5 times by adding 500 μL of bead-binding buffer followed by 5 min incubation on a rotating mixer. The beads with immobilized MpAgo RNP were resuspended in 60 μL of bead-binding buffer. Then the sample was split three ways. Bead-binding buffer, fully complementary target site containing mRNA, or total RNA was added to the three samples, to a final volume of 50 μL for each reaction. After incubation at 37 °C for 1 h, the samples were placed on a magnetic rack. The flow-through fractions were saved for SDS-PAGE analysis. The beads were washed 3 times with 500 μL each wash of bead-binding buffer. To elute MpAgo RNPs, the beads were resuspended in 20 μL of 2× SDS-PAGE loading buffer (100 mM Tris-HCl, 4% (w/v) SDS, 20% (v/v) glycerol, 200 mM DTT, 0.2% (w/v) Bromophenol blue, pH_RT_ 6.8) and boiled at 95 °C for 5 min. The supernatant (elution sample) was separated from the beads using a magnetic rack. The flow-through and elution samples were analyzed by SDS-PAGE.

#### MpAgo Unloading Assay Using Filter-Binding Approach

The MpAgo RNP was prepared by mixing 100 nM of catalytically inactive MpAgo and 0.1 nM of 22 nt long 3′-end radiolabeled gRNA in RNP assembly buffer. The samples were incubated either at 37 °C for 45 min or at 55 °C for 15 min and cooled down slowly to room temperature. 5 μM of either unlabeled fully complementary substrate (ON target) or unlabeled non-complementary ssRNA substrate (OFF target) was added to the MpAgo RNPs in substrate binding buffer (20 mM Tris-HCl, 150 mM KCl, 2 mM MgCl_2_, 5 % (v/v) glycerol, 2 mM DTT, 10 μg/mL BSA, 2 μg/mL heparin, pH_RT_ 7.5). The 50 μL binding reactions were incubated for 1 h at 37 °C. 45 μL of each sample was applied to a dot-blot apparatus with low vacuum and three stacked membranes (Tuffryn (Pall Corporation), Protran (GE Healthcare), Hybond-N^+^ (GE Healthcare)) to separate the free gRNA from gRNA bound to the MpAgo RNP. Membranes were washed with washing buffer, air-dried, and visualized by phosphor-imaging. The intensities of the free gRNA and the gRNA bound to the MpAgo RNP were analyzed using GelEval 1.35 (FrogDance Software).

#### Competition-Binding Assay

The MpAgo-gRNA complex was reconstituted by incubating 300 nM WT MpAgo and 0.1 nM 3′-end radiolabeled gRNA at 55 °C for 15 min in RNP assembly buffer. After adding increasing concentrations of unlabeled 5’-OH or 5’-PO_4_ gRNA, the competition-binding reactions were overlaid with 30 μL of mineral oil (Sigma) to avoid evaporation and were incubated overnight (for 19 hours) at 37 °C. 45 μL of each sample was applied to a dot-blot apparatus with low vacuum and three stacked membranes (Tuffryn (Pall Corporation), Protran (GE Healthcare), Hybond-N^+^ (GE Healthcare)) to separate the MpAgo-gRNA complexes and the free gRNA. The membranes were washed with washing buffer, air-dried, and visualized by phosphor-imaging. The intensities of free and MpAgo-bound gRNA were analyzed using GelEval 1.35 (FrogDance Software). The obtained data of three independent experiments was plotted as a function of competitor (either 5’-OH or 5’-PO_4_ gRNA) concentration, accounting for the ~300 nM MpAgo used in the RNP assembly, and fit with a one-site competitive binding equation using the Prism 6 program (GraphPad Software, Inc.), and the IC_50_ values (i.e. the concentration of competitor needed to displace 50% of the gRNA from the RNP) were determined.

The competition-binding assay to assess the off-rates of gRNAs from MpAgo RNPs reconstituted at 37 °C and 55 °C was performed using the MpAgo-gRNA complexes that were reconstituted by incubating 300 nM WT MpAgo and 0.1 nM 3′-end radiolabeled gRNA either at 37 °C for 45 min or at 55 °C for 15 min. After adding 4 μM of unlabeled 5’-OH or 5’-PO_4_ gRNA, the competition-binding reactions were overlaid with 30 μL of mineral oil (Sigma) to avoid evaporation and were incubated at 37 °C. The 45 μL samples were collected at the various time points ranging from 5 to 2880 min and applied to a dot-blot apparatus with low vacuum and three stacked membranes as described above. The obtained data of three independent experiments were plotted as a function of time and fit with a one phase exponential decay function using the Prism 6 program (GraphPad Software, Inc.), and the average values of the gRNA off-rate (k_off_) and the MpAgo RNP half-life (t_1/2_) were determined.

#### Single Turnover *In Vitro* Cleavage Assays

For the single turnover experiments, the MpAgo-gRNA complexes were reconstituted by mixing 5 nM MpAgo with 5.5 nM gRNA in cleavage buffer (20 mM Tris-HCl, 150 mM KCl, 2 mM MnCl_2_, 5% (v/v) glycerol, 2 mM DTT, pH_RT_ 7.5) and incubating at 55 °C for 15 min. The cleavage reactions were initiated by adding 5’-radiolabeled ssRNA substrates to a final concentration of 0.5 nM and then incubated at 58 °C. 15 μL aliquots were removed at the 0, 0.5, 1, 2, 5, 10, 20, and 30 min time points and quenched by mixing them with an equal volume of formamide gel loading buffer (95% formamide, 100 mM EDTA, 0.025% SDS, and 0.025% (w/v) Bromophenol Blue). Samples were heated to 90 °C for 2 min, resolved on 13% denaturing polyacrylamide gel, and visualized by phosphor-imaging. Assays were performed in three independent replicates, and the intensities of the uncleaved and cleaved RNA were analyzed using GelEval 1.35 (FrogDance Software). The mean ± standard deviation (SD) at each time point of the three independent experiments was plotted as a function of time and the data were fit with an exponential one-phase decay curve using the Prism 6 software (GraphPad Software, Inc.). The first-order cleavage rate constants were determined and reported as apparent k_obs_, because the nature of the rate-limiting step is not known.

#### ssRNA Substrate Filter-Binding Assay

MpAgo-gRNA complexes were reconstituted by incubating catalytically inactive MpAgo and gRNA at 55 °C for 15 min in RNP assembly buffer with a ratio of 1:1.5, if not specified otherwise. The binding reaction was performed in substrate binding buffer (20 mM Tris-HCl, 150 mM KCl, 2 mM MgCl_2_, 5 % (v/v) glycerol, 2 mM DTT, 10 μg/mL BSA, 2 μg/mL heparin, pH_RT_ 7.5) by mixing various concentrations of MpAgo RNP and either 0.01 nM of 5’-radiolabeled complementary ssRNA substrate or non-complementary ssRNA substrate in a final volume of 100 μL to assess MpAgo RNP’s binding affinity and specificity. The binding reactions were incubated for 1 h at 37 °C. 90 μL of each sample was applied to a dot-blot apparatus with low vacuum and three stacked membranes (Tuffryn (Pall Corporation), Protran (GE Healthcare), Hybond-N^+^ (GE Healthcare)) to separate the free ssRNA substrate and the ssRNA substrate bound to the MpAgo RNP. Membranes were washed with washing buffer, air-dried, and visualized by phosphor-imaging. The intensities of the free ssRNA and the ssRNA bound to the MpAgo RNP were analyzed using GelEval 1.35 (FrogDance Software).

#### Crosslinking of 5’-end BrdU gRNA to MpAgo

The MpAgo-gRNA complex was prepared by mixing MpAgo and guide RNA that contains 5-bromo-2′-deoxyuridine at the 5’-terminal 1^st^ position with a ratio of 1:1.5, or 10 μM MpAgo and 15 μM 5’-BrdU gRNA (if not specified otherwise) in RNP assembly buffer described above. After mixing the components, the sample was incubated at 55 °C for 15 min and cooled down slowly to room temperature. Next, crosslinking was performed by irradiating reconstituted MpAgo RNP in an uncapped 0.5 mL tube at ~ 305 nm for 1 h on ice (using Hg-UVB lamp bulbs and Spectrolinker XL-1500 UV Crosslinker (Spectronics Corporation Spectroline)). The efficiency of crosslinking was assessed using 200 pM 3′-end radio labeled gRNAs incubated with 5 μM MpAgo. After the crosslinking the MpAgo RNPs were resolved on gradient 4 – 20 % SDS-PA gel, and visualized by phosphor-imaging. Assays were performed in three independent replicates and the intensities of the crosslinked and non-crosslinked gRNA were analyzed using GelEval 1.35 (FrogDance Software). The mean ± standard deviation (SD) for each gRNA was graphed in a bar plot.

#### Optimization of ssRNA binding assay

The 5’-end modification of gRNA improved ON target substrate binding specificity and affinity (Figs. 2C, S5) and stabilized the MpAgo RNP (Fig. S5A), when using MpAgo RNP reconstituted at 1:1.5 MpAgo:gRNA ratio and 2 μg/mL of heparin. The further increases in the protein:gRNA ratio did not have a significant effect on the MpAgo RNP binding amplitude of the ON and OFF target RNAs (Fig. S6A). We therefore used the 1:1.5 ratio of MpAgo:gRNA for reconstitution and crosslinking in all further experiments.

The ON target binding amplitude was slightly improved while non-specific OFF target binding was reduced by supplementing the reaction with 10 μg/mL of heparin (Fig. S6B). Further increases in heparin concentration diminished not only OFF target binding but also ON target binding (Fig. S6B). Thus, we used 10 μg/mL of heparin in subsequent experiments to assess ssRNA binding.

The optimal MpAgo RNP concentration for binding specificity and efficiency was determined by ssRNA substrate filter-binding assays performed at 37 °C, with 0.1 nM of either fully complementary substrate or non-complementary substrate in the presence of 10 μg/mL heparin. The lowest concentration of the RNP tested (10 nM) provided the most specific binding (Fig. S6C).

To obtain MpAgo RNP – substrate RNA equilibrium binding isotherms, 10 pM of 5’-radiolabeled RNA substrate was incubated with increasing concentrations of preassembled MpAgo RNP in a final volume of 100 μL. The binding reactions were incubated for 1 h at 37 °C. 95 μL of each sample was applied to a dot-blot apparatus with low vacuum and three stacked membranes as described above. The membranes were washed with washing buffer, air-dried, and visualized by phosphor-imaging. The intensities of the free ssRNA and the ssRNA bound to the MpAgo RNP were analyzed using GelEval 1.35 (FrogDance Software).

#### Isolation of A-to-I Edited ssRNA Substrates from a Complex Mixture

The MpAgo-gRNA complex was reconstituted by incubating catalytically inactive MpAgo and gRNA at 55 °C for 15 min in the RNP assembly buffer with a ratio of 1:1.5. Next, crosslinking was performed by incubating reconstituted MpAgo RNP under UVB light (λ_max_ ~ 305 nm) for 1 h on ice. The MpAgo RNP binding reaction was performed in substrate binding buffer (20 mM Tris-HCl, 150 mM KCl, 2 mM MgCl_2_, 5 % (v/v) glycerol, 2 mM DTT, 10 μg/mL BSA, pH_RT_ 7.5) by mixing 0.5 nM of the MpAgo RNP and 0.1 nM of 5’-radiolabeled ssRNA target containing either I or A, in the presence of 500 ng of total RNA purified from HEK 293T cells and 200 ng/μL of yeast tRNA (total reaction volume 50 μL). The binding reactions were incubated for 1 h at 37 °C. 45 μL of each sample was applied to a dot-blot apparatus with low vacuum and three stacked membranes (Tuffryn (Pall Corporation), Protran (GE Healthcare), Hybond-N^+^ (GE Healthcare)) to separate the free ssRNA substrate and the ssRNA substrate bound to the MpAgo RNP. Membranes were washed with washing buffer, air-dried, and visualized by phosphor-imaging. The intensities of the free ssRNA and the ssRNA bound to the MpAgo RNP were analyzed using GelEval 1.35 (FrogDance Software).

---

**Figure S1.**
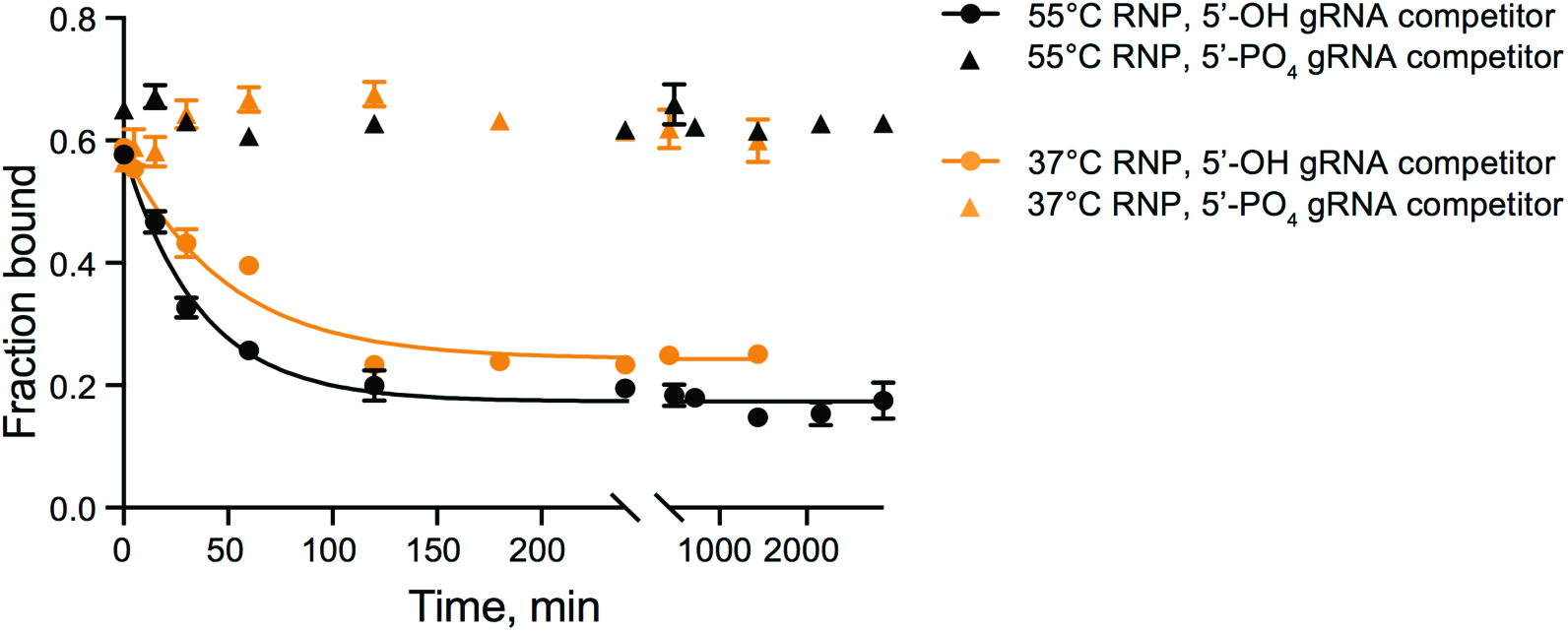
Stability of the *in vitro* reconstituted MpAgo-gRNA complex. Stability of MpAgo RNPs at 37 °C was assessed by measuring the displacement of gRNA from the RNPs reconstituted at 37 °C (orange) and 55°C (black) using 300 nM MpAgo and 0.1 nM 3′-end radiolabeled gRNA after the addition of 4 μM 5’-OH RNA competitor (circles) and μM 5’-PO_4_ RNA competitor (triangles). The obtained average off-rate (k_off_) of the 5’-OH gRNA pre-bound to MpAgo at 55 °C is 0.027 ± 0.002 min^−1^, and k_off_ of the 5’-OH gRNA pre-bound to MpAgo at 37 °C is 0.021 ± 0.002 min^−1^. The half-life time (t_1/2_) of the RNP assembled at 55°C is ~25 min, while the t_1/2_ of the RNP assembled at 37°C is ~34 min. Data are represented as the mean ± SD from three independent experiments.

**Figure S2.**
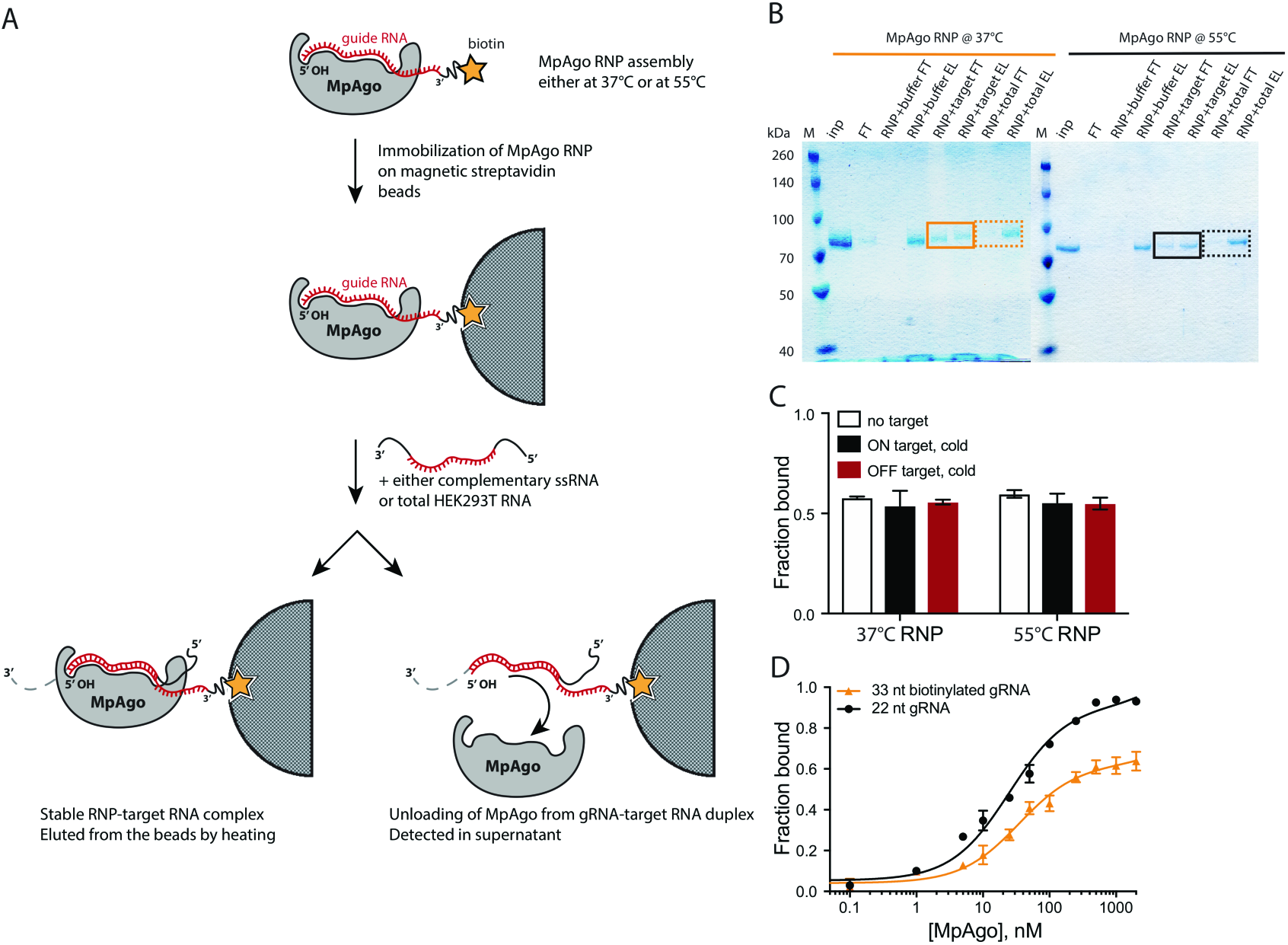
Stability of the *in vitro* reconstituted MpAgo-gRNA complex upon complementary substrate RNA binding. **(A)** A scheme depicting the MpAgo unloading assay. MpAgo RNP was assembled either at 37 °C or at 55 °C, using catalytically inactive MpAgo and biotinylated 33-nucleotide guide RNA, and then immobilized on streptavidin-coated magnetic beads. The beads were washed 5× with bead-washing buffer to remove unbound RNPs. Next, the immobilized RNPs were incubated with only the buffer, the fully complementary substrate RNA (ON), or a pool of non-complementary RNAs (OFF). The flow-through fractions were saved for SDS-PAGE analysis. The beads were washed 3 times with bead-washing buffer. MpAgo RNPs bound to the beads were eluted by boiling. The flow-through and elution samples were analyzed by 12% SDS-PAGE. **(B)** SDS gel of MpAgo bound to and released from the gRNA upon adding the complementary ssRNA substrate. The MpAgo unloading assay was performed to test the temperature at which the assembled RNP is more stable in the presence of the ssRNA substrate. The collected flow-through (FT) and elution (E) fractions were resolved on 12% SDS-PAGE. MpAgo RNPs assembled at 37 °C (solid, orange box); MpAgo RNPs assembled at 55 °C (solid, black box). No MpAgo protein was detected in the FT fraction after adding non-specific RNAs (dashed, orange and black boxes). **(C)** The stability of the MpAgo RNP complexes (assembled at two different temperatures using 3′-end radiolabeled 22-nt gRNA) upon the interaction with RNA substrate assessed by filter-binding. The reconstituted RNPs (300 nM MpAgo with 0.1 nM gRNA) were incubated for 1 h at 37 °C with only the buffer (white), 5 μM of the fully complementary substrate RNA (ON target, black), or 5 μM of the non-complementary RNA (OFF target, red). The average fraction of bound gRNA is represented in a bar plot. **(D)** The MpAgo and gRNA binding curves obtained for the 22-nucleotide (black) and 33-nucleotide (orange) gRNAs at 55 °C. The fraction of the bound gRNAs is plotted as a function of the MpAgo concentration. The data were fit with a standard binding isotherm (solid lines) to extract the binding affinities. MpAgo binds the 22-nucleotide and 33-nucleotide gRNAs with K_d_ values of 25 ± 3 nM and 35 ± 7 nM, respectively. Data are represented as the mean ± standard deviation (SD) from three independent experiments.

**Figure S3.**
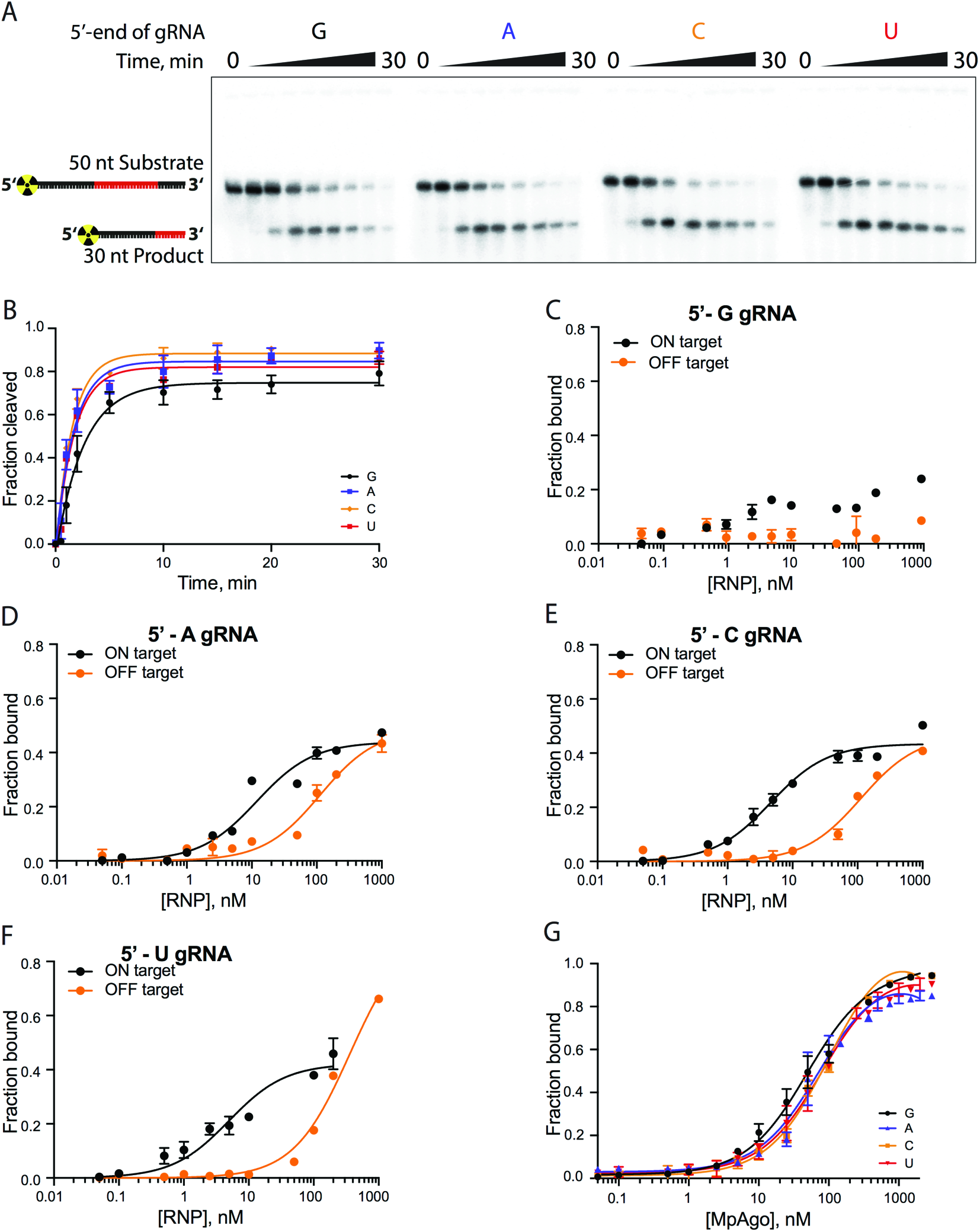
The role of the 5’-terminal nucleotide of the gRNA in substrate ssRNA cleavage and binding by MpAgo RNP. **(A)** A representative polyacrylamide denaturing gel of the 5’-radiolabeled ssRNA substrate cleavage reactions performed with MpAgo RNP programmed with a gRNA containing 5’-terminal guanosine (G), adenosine (A), cytidine (C) or uridine (U). **(B)** The fraction of cleaved ssRNA substrate plotted as a function of time. The data were fit with a single exponential to extract the apparent cleavage rate constants (k_obs_): 5’-G k_obs_ = 0.39 ± 0.05 min^−1^; 5’-A k_obs_ = 0.59 ± 0.09 min^−1^; 5’-U k_obs_ = 0.59 ± 0.07 min^−1^; and 5’-C k_obs_ = 0.64 ± 0.09 min^−1^. Data are represented as the mean ± SD from three independent experiments. **(C-F)** The ssRNA filter-binding assay performed with MpAgo RNP programmed with the 5’-G, 5’-A, 5’-U, or 5’-C guide RNA and fully complementary (21-nucleotide complementarity, ON target, black) or non-complementary RNA substrates (OFF target, orange). The fraction of the bound ssRNA substrate was plotted as a function of the MpAgo RNP concentration. For 5’-G gRNAs the K_d_ was not determined due to poor binding of the ON target to the MpAgo RNP **(C)**. The obtained average K_d_ for ON and OFF target RNAs of the MpAgo RNP programmed with 5’-A gRNA are 12 ± 2 nM and 112 ± 19 nM respectively **(D)**, with 5’-C gRNA, 4.6 ± 0.5 nM and 112 ± 18 nM respectively **(E)**, and with 5’-U gRNA, 5.0 ± 0.8 nM and 359 ± 37 nM respectively **(F)**. Data are represented as the mean ± SD from three independent experiments. **(G)** Binding curves of the MpAgo and the 5’-G, 5’-A, 5’-U, or 5’-C guide RNA obtained from filter binding assays at 55 °C. The fraction of bound gRNA was plotted as a function of the MpAgo concentration. The obtained average K_d_’s for MpAgo and the gRNAs with 5’-guanine, adenine, uracil or cytosine are: 5’-G K_d_ = 49 ± 5 nM; 5’-A K_d_ = 73 ± 10 nM; 5’-U K_d_ = 81 ± 9 nM; and 5’-C K_d_ = 104 ± 9 nM. Data are represented as the mean ± standard deviation (SD) from three independent experiments.

**Figure S4.**
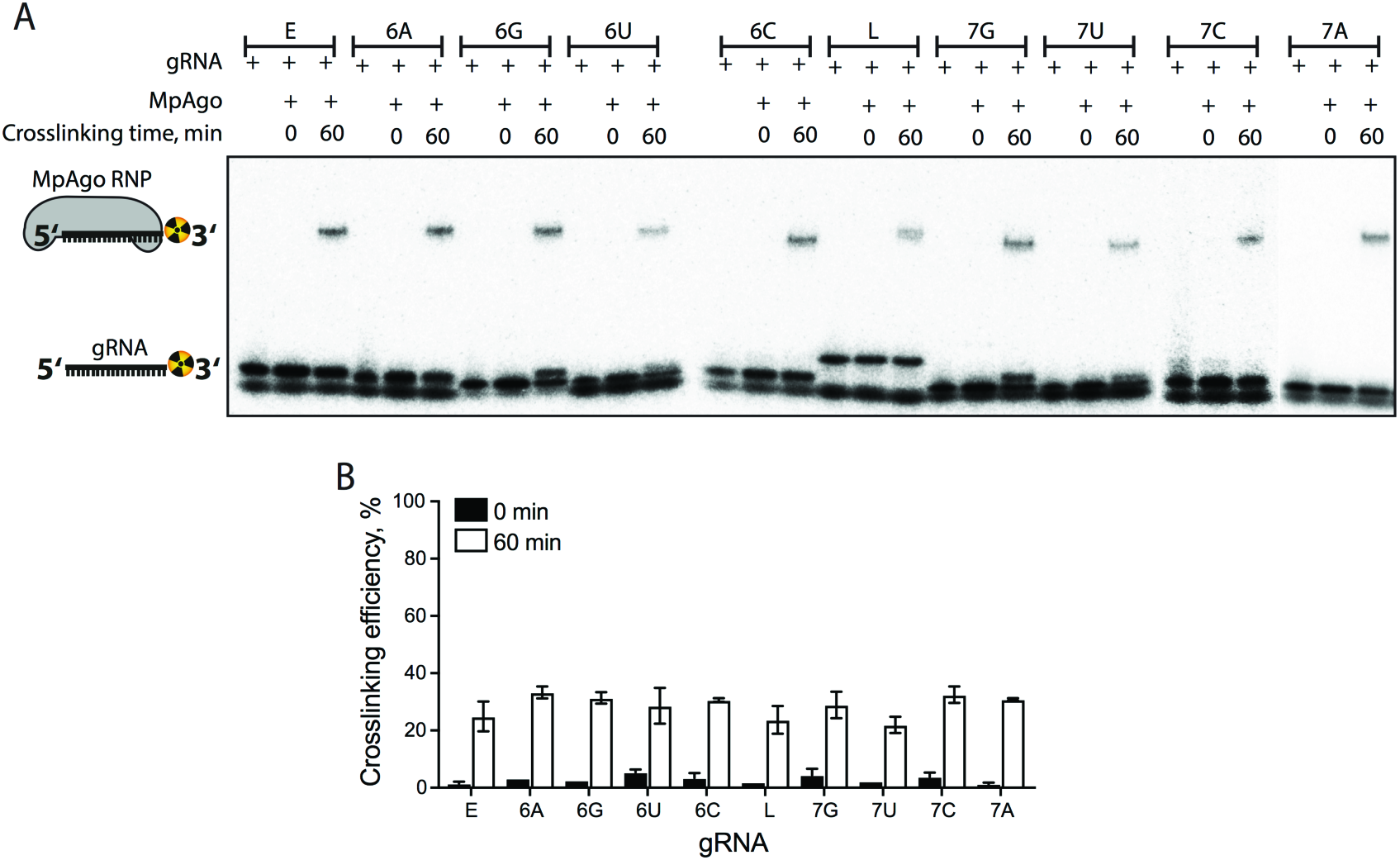
The efficiency of crosslinking 5’-BrdU gRNA to MpAgo. **(A)** A representative 4 - 20 % gradient SDS gel of the 3′-end radiolabeled gRNA crosslinking reactions was performed with MpAgo RNPs programmed with the 5’-end BrdU gRNAs used in the binding and cleavage experiments (**Table S1**). The lowest band is unincorporated radiolabel. **(B)** The fraction of crosslinked gRNA represented in a bar plot. Data are represented as the mean ± standard deviation (SD) from three independent experiments.

**Figure S5.**
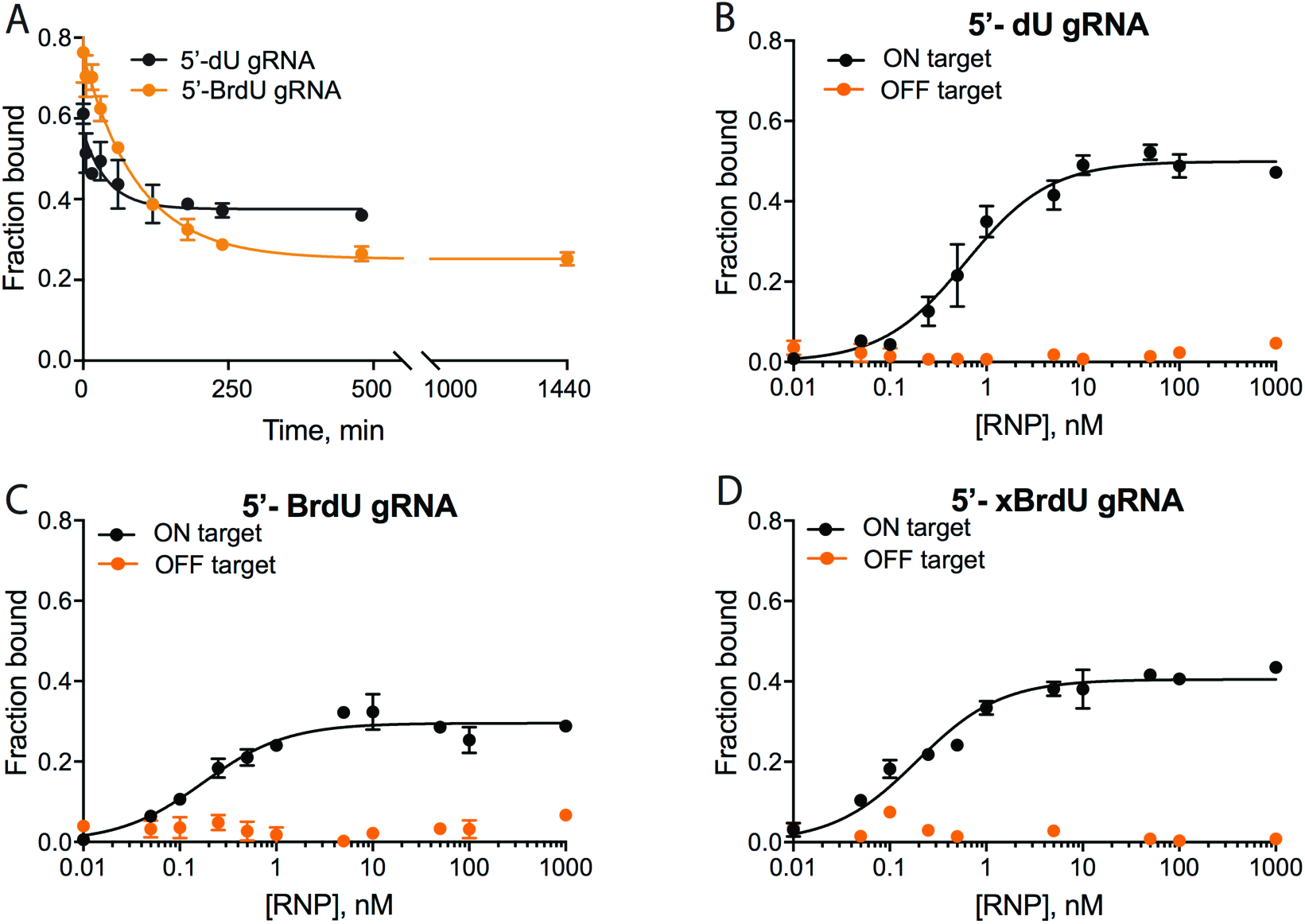
Stability, affinity, and specificity of the MpAgo RNPs programmed with 5’-end modified gRNAs. **(A)** Stability of the MpAgo RNPs at 37 °C measured by displacement of gRNA from the RNPs programmed at 55 °C using 300 nM MpAgo and 0.1 nM 5’-dU (orange) and 5’-BrdU (black) gRNAs, after the addition of 4 μM cold gRNA competitor. The obtained average off-rates (k_off_) of 5’-dU and 5’-BrdU gRNAs pre-bound to MpAgo at 55 °C are 0.024 ± 0.008 min^−1^, and 0.011 ± 0.001 min^−1^, respectively. The half-life times (t_1/2_) of the RNPs programmed with 5’-dU and 5’-BrdU gRNAs are ~29 min and ~66 min, respectively. Data are represented as the mean ± SD from three independent experiments. **(B-D)** The ssRNA filter-binding assay was performed using MpAgo RNP programmed with 5’-dU **(B)** or 5’-BrdU **(C)** gRNAs or with a BrdU gRNA with subsequent 1h exposure to 305 nm UV light for crosslinking (5’-xBrdU) **(D)**. The obtained average K_d_ values of these MpAgo RNPs for the fully complementary substrate RNA (ON target, black) are: 5’-dU K_d_ = 500 ± 67 pM **(B)**; 5’-BrdU K_d_ = 175 ± 23 pM **(C)**; and 5’-xBrdU K_d_ = 193 ± 20 pM **(D)**. The K_d_ values of these MpAgo RNPs for the non-complementary RNA substrates (OFF target, orange) were not determined due to undetectable binding. Data are represented as the mean ± standard deviation (SD) from three independent experiments.

**S6.**
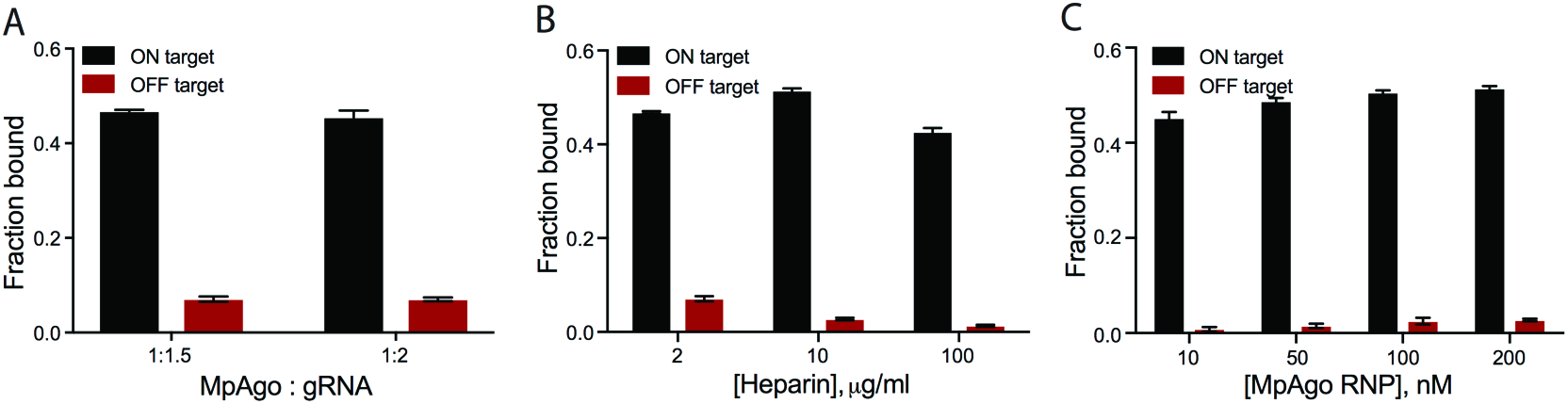
ssRNA substrate binding conditions using MpAgo RNP loaded with 5’-BrdU gRNA. **(A)** Optimal MpAgo:gRNA ratio for the reconstitution of the MpAgo RNP programmed with 5’-BrdU modified gRNA was determined using ssRNA substrate filter-binding assays performed at 37 °C, where 200 nM of the preassembled MpAgo RNPs were incubated with either fully complementary substrate (ON target, black) or non-complementary substrate (OFF target, red) at 0.1 nM concentration. The fraction of the ssRNA substrate bound to the RNP was quantified and plotted as the mean ± SD from three independent experiments. **(B)** The optimal heparin concentration in the ssRNA binding reaction was determined by ssRNA substrate filter-binding assays performed at 37 °C, with either 0.1 nM of fully complementary substrate (ON target, black) or non-complementary substrate (OFF target, red) and 200 nM MpAgo RNP at three heparin concentrations: 2 μg/mL, 10 μg/mL and 100 μg/mL. The fraction of the ssRNA substrate bound to the RNP was quantified and plotted as the mean ± SD from three independent experiments. **(C)** The optimal MpAgo RNP concentration was determined by ssRNA substrate filter-binding assays performed at 37 °C, with either 0.1 nM of fully complementary substrate (ON target, black) or non-complementary substrate (OFF target, red) in the presence of 10 μg/mL heparin at four MpAgo RNP concentrations. The fraction of the ssRNA substrate bound to the RNP was quantified and plotted as the mean ± SD from three independent experiments.

**Figure S7.**
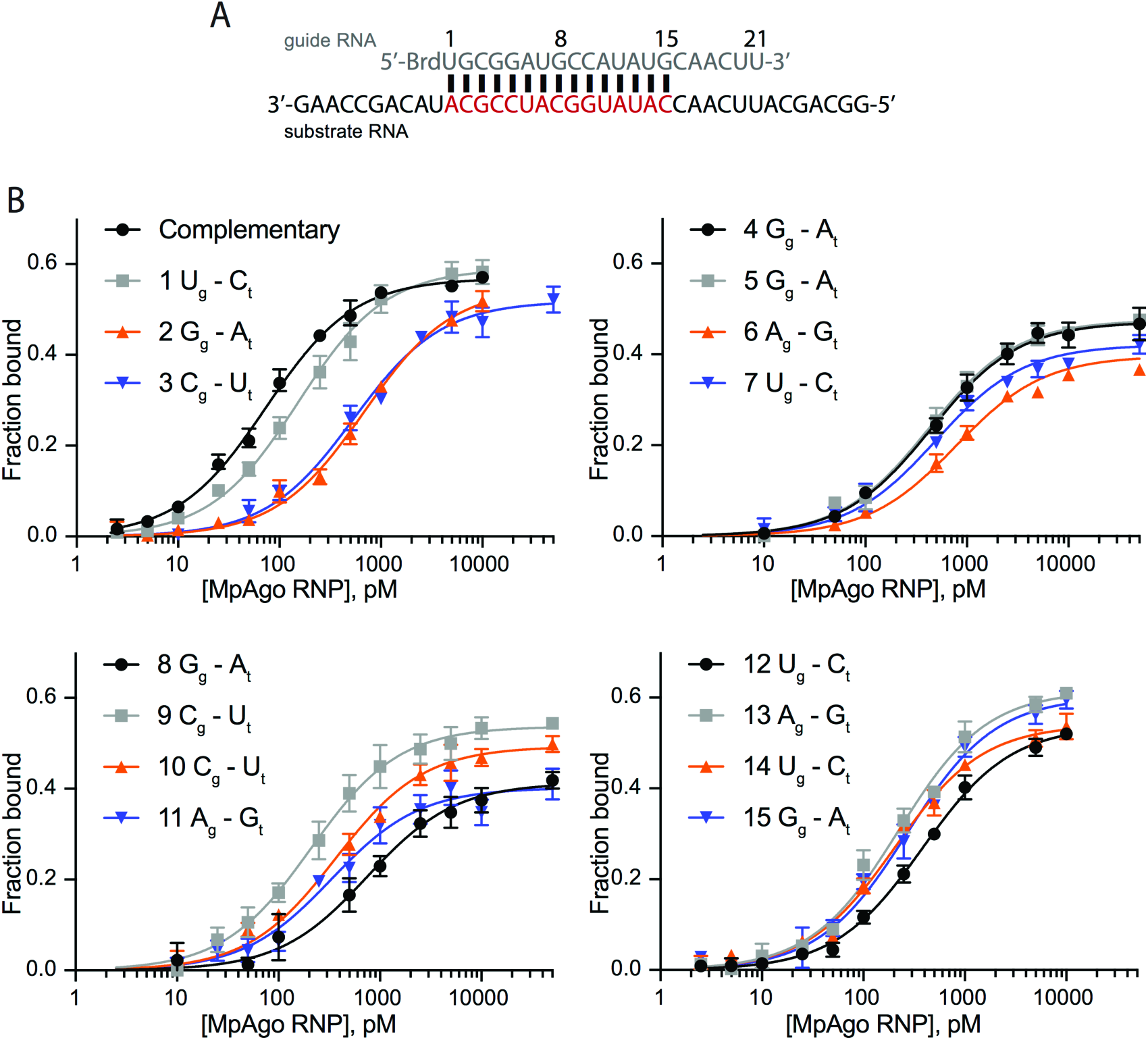
Binding of 5’-BrdU MpAgo RNP and the ssRNA substrates containing single mismatches across the gRNA. **(A)** A schematic depiction of the sequences of the 5-bromo-deoxyuridine modified gRNA (grey) and the complementary ssRNA substrate (black and complementary region in red) used for binding assays. Single nucleotide mismatches were introduced across the red region keeping the sequence of the gRNA constant. **(B)** The binding curves of MpAgo RNP and the ssRNA substrates containing mismatches at various positions from the 1^st^ to the 15^th^ position. The position and nature of the mismatches are indicated in the legends of the plots, where a nucleotide with a subscript ‘g’ is located in the gRNA and a nucleotide with a subscript ‘t’ is located in the ssRNA target strand. The data were fit with a standard binding isotherm (solid lines) to extract the binding affinities. The ratios of obtained average dissociation constants for the mismatches (K_d_^mm^) to the fully complementary substrate (K_d_^Compl^) were calculated and plotted against the position of the mismatch in Fig. 4A. The ratios of the obtained average dissociation constants for the purine-purine mismatches (K_d_^mm^) to the fully complementary substrate (K_d_^Compl^) were calculated and plotted against the position (1^st^, 2^nd^, 4^th^, 5^th^, 6^th^, 8^th^, 11^th^, 13^th^, and 15^th^) of the mismatch in Fig. 4B.

**Figure S8.**
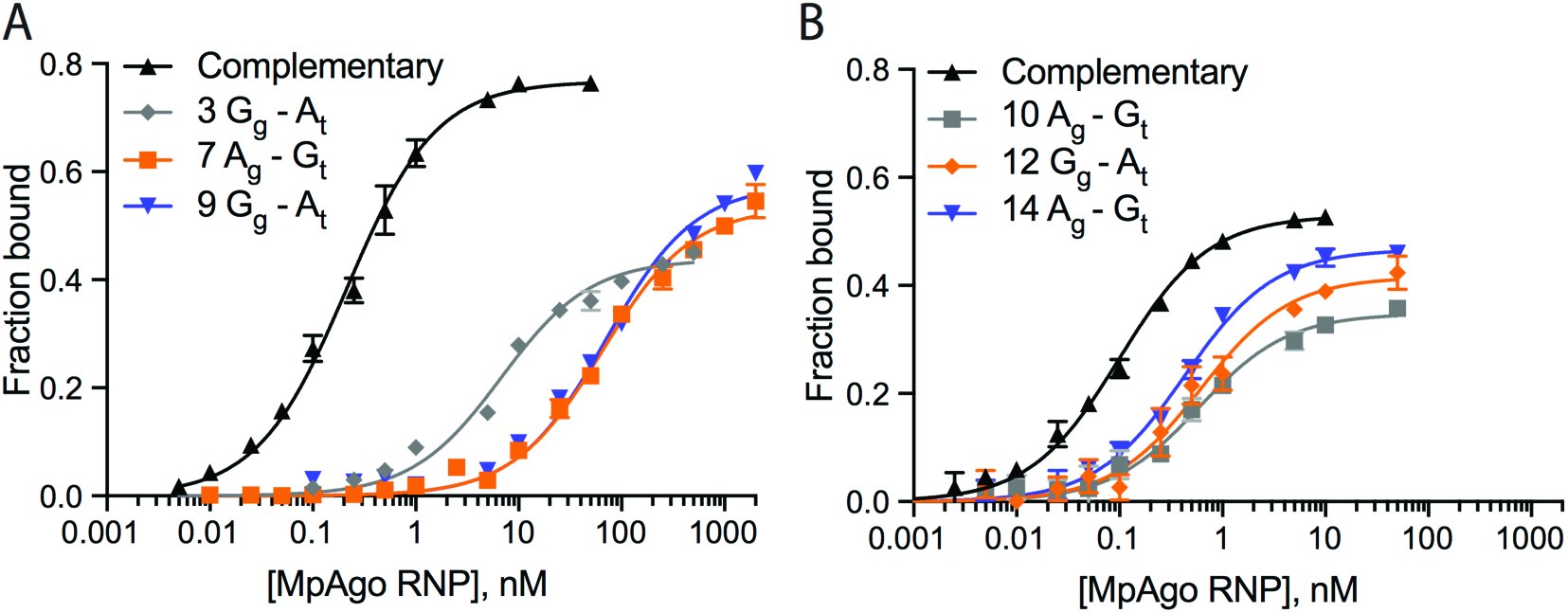
Binding of 5’-BrdU MpAgo RNPs and ssRNA substrates containing purine-purine mismatches. To obtain purine-purine mismatches at all 15 positions, we used two additional gRNAs (**Table S1**) to program MpAgo. **(A)** Binding curves of MpAgo RNP and ssRNA substrates that create purine-purine mismatches at the 3^rd^, 7^th^ and 9^th^ positions of the gRNA. **(B)** Binding curves of MpAgo RNP and ssRNA substrates that create purine-purine mismatches at the 10^th^, 12^th^ and 14^th^ positions of the gRNA. The position and nature of the mismatches are indicated in the legends of the plots, where a nucleotide with a subscript ‘g’ is located in the gRNA and a nucleotide with a subscript ‘t’ is located in the ssRNA target strand. The data were fit with a standard binding isotherm (solid lines) to extract the binding affinities. The ratios of the obtained average dissociation constants for the purine-purine mismatches (K_d_^mm^) to the fully complementary substrate (K_d_^Compl^) were calculated and plotted against the position of the mismatch in Fig. 4B.

**Figure S9.**
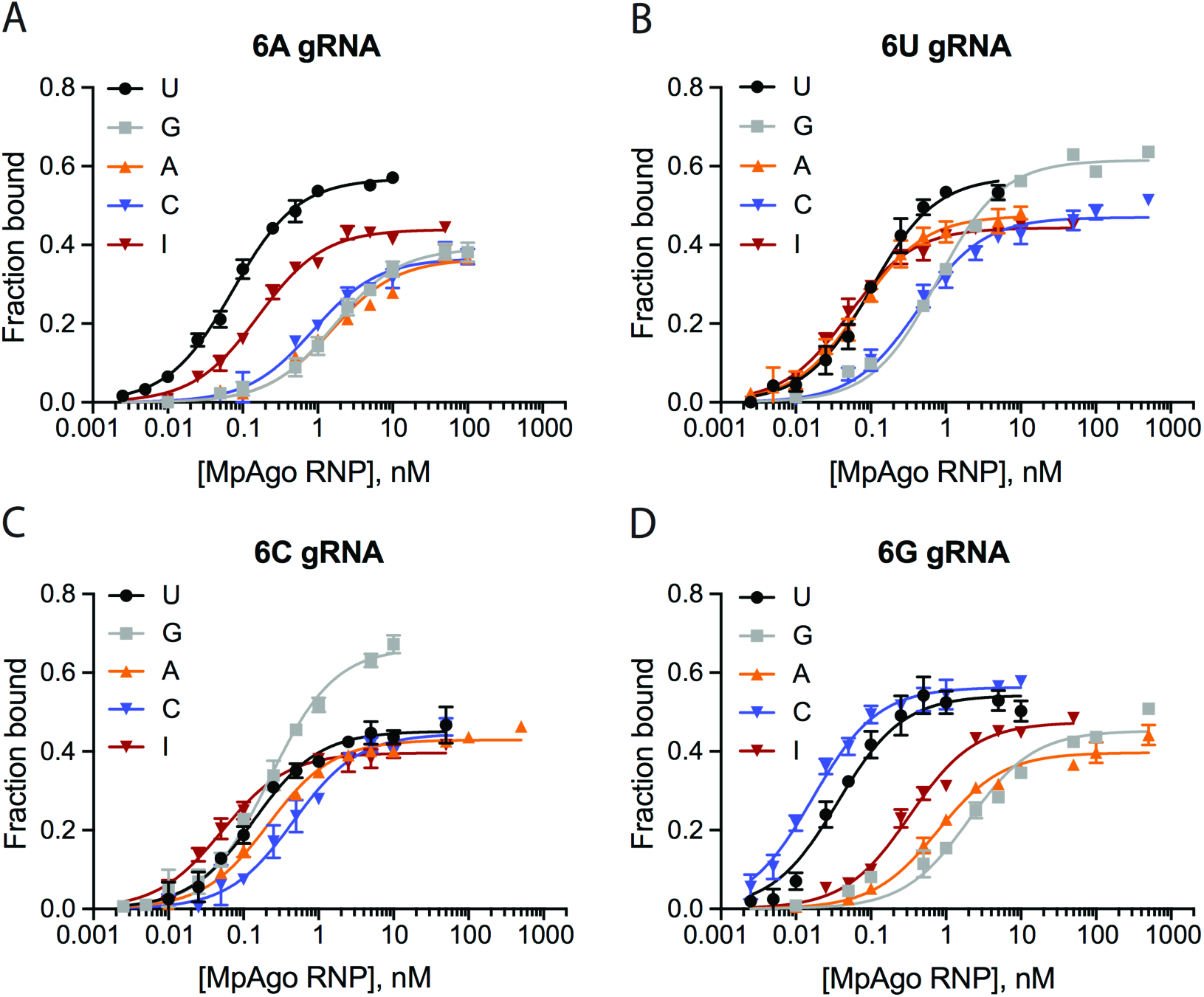
Binding of 5’-BrdU MpAgo RNP programmed with gRNAs containing permutations at the 6^th^ position and ssRNA substrates containing permutations at the corresponding position. The ssRNA filter-binding assays were performed using MpAgo RNPs programmed with A **(A)**, U **(B)**, C **(C)**, or G **(D)** gRNA at the 6^th^ position and ssRNA targets containing U, C, A, G or I at the corresponding position. The average K_d_’s were extracted and are represented in a heat map (Fig. 5A).

**Figure S10.**
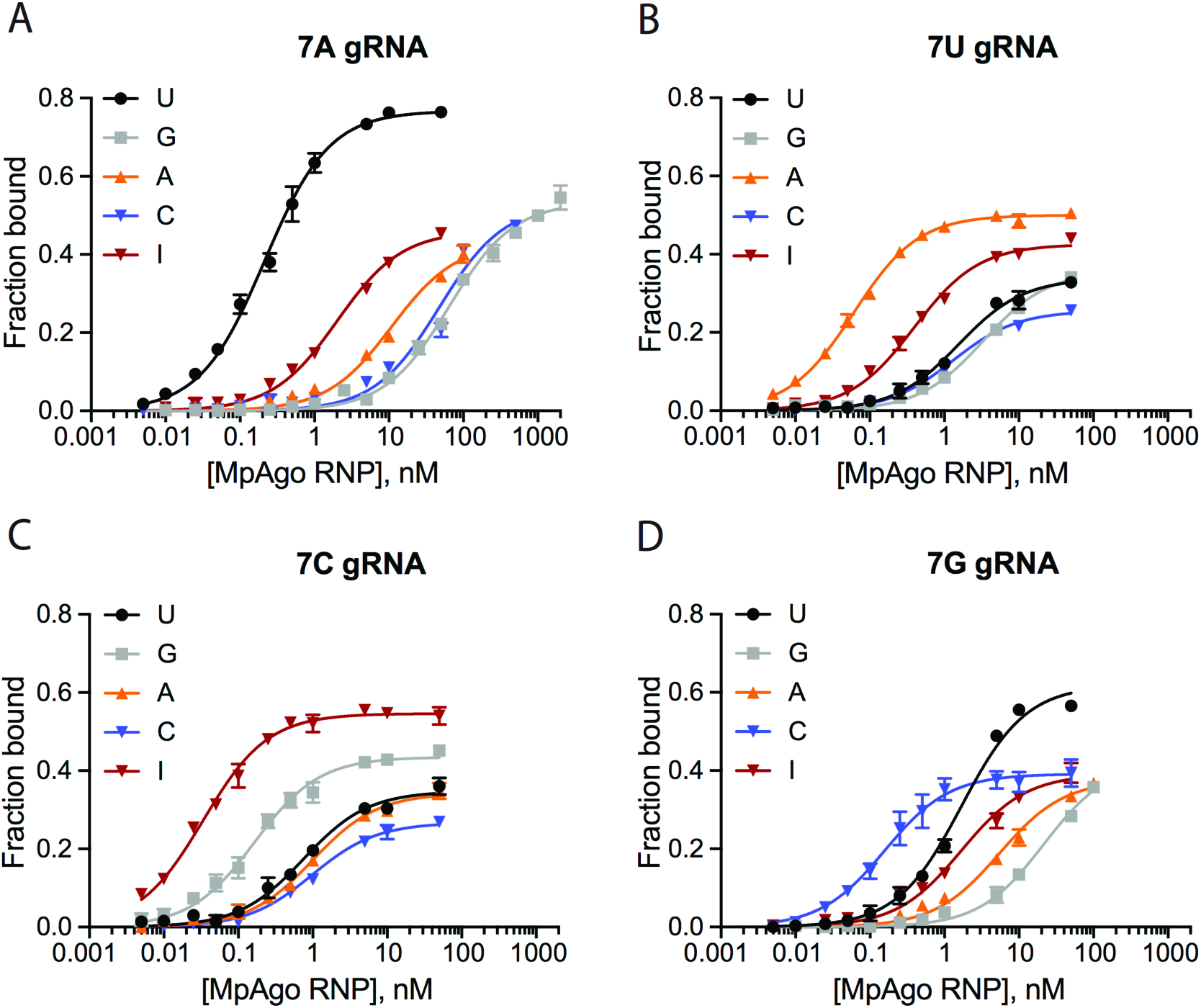
Binding of 5’-BrdU MpAgo RNP programmed with gRNAs containing permutations at the 7^th^ position and ssRNA substrates containing permutations at the corresponding position. The ssRNA filter-binding assays were performed using MpAgo RNPs programmed with A **(A)**, U **(B)**, C **(C)**, or G **(D)** gRNA at the 7^th^ position and ssRNA targets containing U, C, A, G or I at the corresponding position. The average K_d_’s were extracted and represented in a heat map (Fig. 5B).

**Figure S11.**
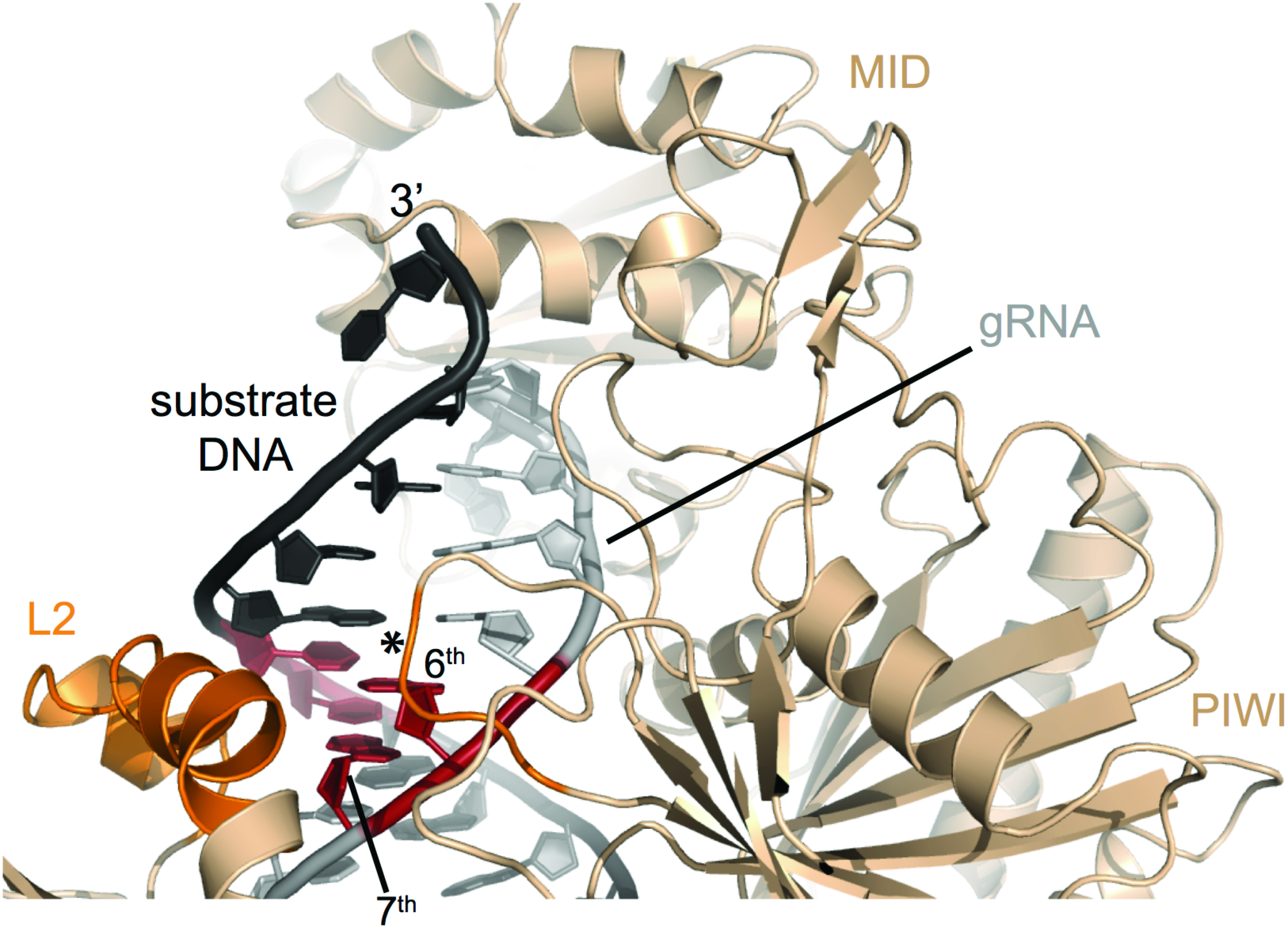
Structure of MpAgo RNP bound to a ssDNA substrate. MpAgo (light orange) bounds to its gRNA (grey) and a fully complementary ssDNA substrate (black). Two helices of the L2 domain and a loop (asterisk) of the PIWI domain (dark orange) probe the shape of the minor groove of the gRNA-ssDNA duplex at the 6^th^ and 7^th^ position (red) of the gRNA. Structural model derived from PDB coordinates 5UX0 (4).

**Table S1.**
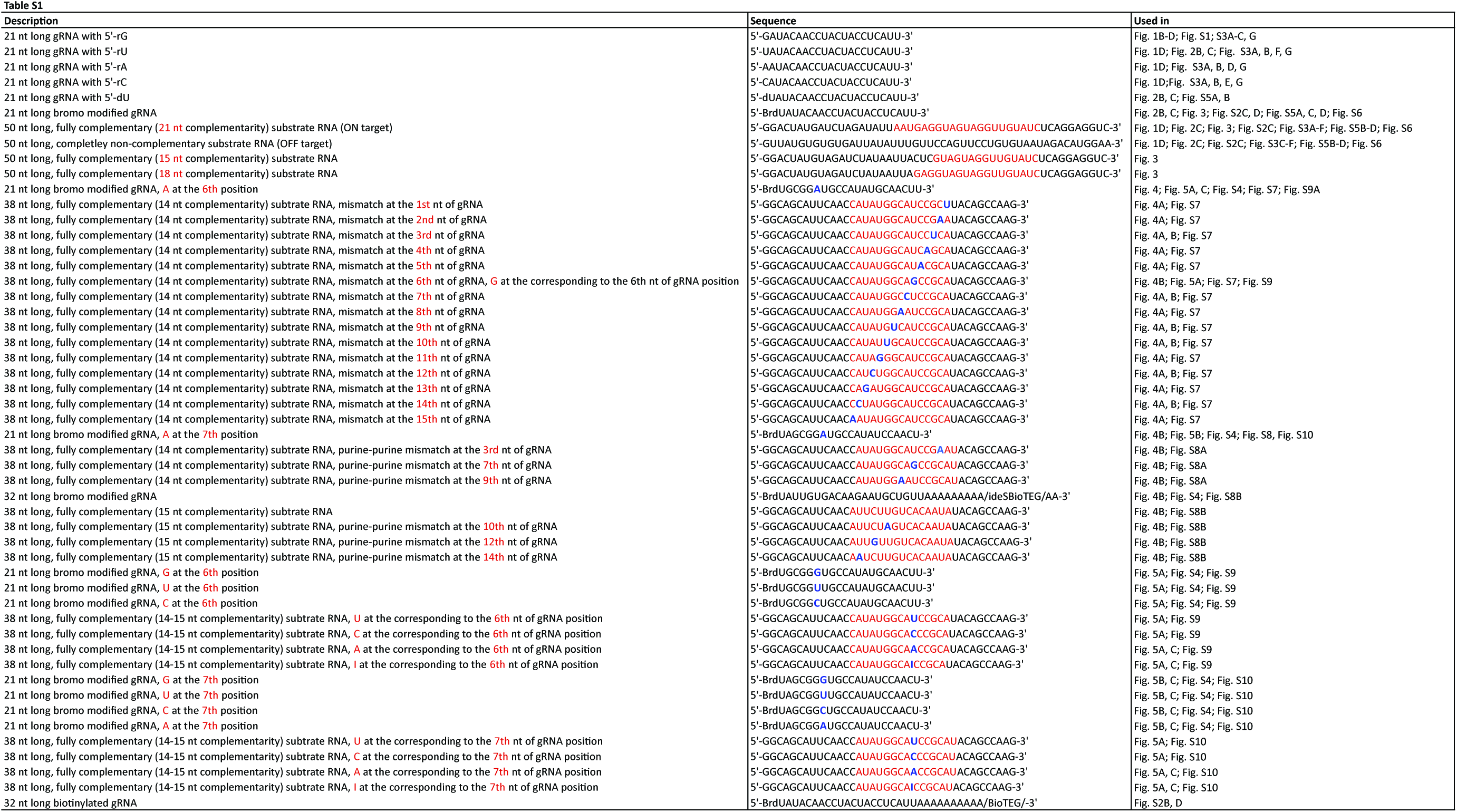

